# Systems biology illuminates alternative metabolic niches in the human gut microbiome

**DOI:** 10.1101/2022.09.19.508335

**Authors:** Cecilia Noecker, Juan Sanchez, Jordan E. Bisanz, Veronica Escalante, Margaret Alexander, Kai Trepka, Almut Heinken, Yuanyuan Liu, Dylan Dodd, Ines Thiele, Brian DeFelice, Peter J. Turnbaugh

## Abstract

Human gut bacteria perform diverse metabolic functions with consequences for host health. The prevalent and disease-linked Actinobacterium *Eggerthella lenta* performs several unusual chemical transformations, but it does not metabolize sugars and its core growth strategy remains unclear. To obtain a comprehensive view of the metabolic network of *E. lenta*, we generated several complementary resources: defined culture media, metabolomics profiles of strain isolates, and a curated genome-scale metabolic reconstruction. Stable isotope-resolved metabolomics revealed that *E. lenta* uses acetate as a key carbon source while catabolizing arginine to generate ATP, traits which could be recapitulated *in silico* by our updated metabolic model. We compared these *in vitro* findings with metabolite shifts observed in *E. lenta-*colonized gnotobiotic mice, identifying shared signatures across environments and highlighting catabolism of the host signaling metabolite agmatine as an alternative energy pathway. Together, our results elucidate a distinctive metabolic niche filled by *E. lenta* in the gut ecosystem.

## INTRODUCTION

Human gut bacteria perform diverse and specialized metabolic functions with consequences for host health. Yet the core metabolic strategies relied upon for growth by many commensal gut microbes remain unclear, which is reflected in the large number of gut taxa that remain difficult to culture (Lagkouvardos et al., 2017; Tramontano et al., 2018). The growth strategies of individual gut species and strains shape their ability to colonize a host and their potential chemical interactions with other community members and with the host (Alexander et al., 2021; Medlock et al., 2018). Efforts to describe and model the metabolism and growth of various community members have included detailed biochemical studies of resource utilization by individual model species such as members of the genus *Bacteroides* (Koropatkin et al., 2012) and *Clostridium sporogenes* (Liu et al., 2022), as well as large-scale efforts to characterize species-level metabolic activity using community multi-omic profiling (Franzosa et al., 2018; Hertel et al., 2019). However, these efforts have been most fruitful for members of the microbiota that are found at high abundance and with prior knowledge of well-annotated metabolic pathways.

One key group of human gut microbes whose core metabolism remains particularly unclear are those that are fully asaccharolytic; *i.e.* derive no growth benefit from sugars and instead may rely on a range of more unconventional nutrients. Many of these taxa are members of the family *Eggerthellaceae*, which are widely found in mammalian gut microbiota (Almeida et al., 2019) but rarely found in other environments. The species *Eggerthella lenta* is a notable example of this group. *E. lenta* is a gram-positive facultative anaerobe found at high prevalence in human gut microbiota (Koppel et al., 2018). Although *E. lenta* is commonly found in healthy individuals, it can cause severe bacteremia (Gardiner et al., 2015) and is increased in abundance in the gut microbiota of patients with several autoimmune diseases (Cekanaviciute et al., 2017; Chen et al., 2016; Islam et al., 2021; Zhu et al., 2021).

*E. lenta* has distinctive metabolic properties and a capacity for many unusual chemical transformations, but it remains unknown how these properties fit into its overall metabolic network and evolutionary strategy. *E. lenta* strains can metabolize varied mammalian and dietary substrates, including cardenolides, bile acids, plant lignans, and dopamine (Bess et al., 2020; Devlin and Fischbach, 2015; Haiser et al., 2013; Koppel et al., 2018; Maini Rekdal et al., 2019). However, none of these compounds except dopamine have been reported to provide a growth or fitness advantage in any conditions tested to date. Genome analysis of *E. lenta* has also predicted that it may be able to perform autotrophic acetogenesis (Harris et al., 2018), but this prediction has not been biochemically validated. *E. lenta* culture conditions typically require rich media and high levels of the amino acid L-arginine. Past studies reported little to no growth of *E. lenta* in minimal or chemically defined media formulations (Hylemon et al., 2018; Maini Rekdal et al., 2020; Tramontano et al., 2018), complicating mechanistic biochemical studies of its metabolism.

In this study, we first developed a chemically defined media that supports strong growth of *E. lenta* strains and described the metabolic footprint and growth determinants of *E. lenta* in this environment. We used stable isotope-resolved metabolomics (SIRM) to investigate the pathways by which *E. lenta* metabolizes two key nutrients, acetate and arginine. This platform allowed us to curate and interpret a genome-scale metabolic model of the *E. lenta* type strain to make predictions about untested growth conditions and to identify gaps in the metabolic network representing novel enzymes or pathways. Extending this approach, we further documented extensive diversity in the metabolic footprint of a collection of *E. lenta* strain isolates. Finally, we evaluated the relevance of these findings to a host-associated context by profiling the metabolome of *E. lenta*-colonized gnotobiotic mice, defining shared and divergent metabolic activities between *in vitro* and *in vivo* environments. In total, we elucidate an unusual metabolic niche and lay a comprehensive foundation for future mechanistic studies of *E. lenta* metabolism.

## RESULTS

### Extensive metabolite footprint of *Eggerthella lenta* in chemically defined media

To identify key nutrients and metabolic pathways required for growth of *E. lenta*, we first developed a custom chemically defined media formulation, referred to as *<u>E</u>ggerthella* <u>D</u>efined Media 1 (EDM1). We designed the initial EDM1 formulation by making several modifications to a recipe previously reported to support growth of many human gut bacterial isolates but not *E. lenta* (Tramontano et al., 2018). We increased the quantity of L-arginine, removed sugars, and ensured the availability of all amino acids and vitamins/cofactors with fragmented or missing biosynthetic pathways in the *E. lenta* DSM 2243 genome [Virtual Metabolic Human database annotations (Noronha et al., 2018), *Methods*, **Table S1**]. The resulting media is composed of compounds typically present in the mammalian gut from microbial, host, and/or dietary sources. It supported robust *E. lenta* growth at a level comparable with standard culture conditions (Brain Heart Infusion media supplemented with 1% arginine; **Figure S1A-B**).

Using this platform, we sought to identify primary metabolites used and produced by *E. lenta*, and the underlying core metabolic pathways active in the EDM1 condition. We used untargeted metabolomics to analyze culture supernatants of the type strain *E. lenta* DSM 2243 across 6 time points over its 50-hour growth curve in EDM1 batch culture (Figure 1A). After dereplication of features from positive and negative ionization modes, 4,095 features were detected, of which 636 (15.6%) were not detected in sterile control media (sample mean intensity > 3x blank sample mean, Figure 1B). 612 features (14.9% of features overall) were significantly different in abundance between sterile controls and supernatants at the final time point (FDR-adjusted *p*<0.1, Figure 1C), of which the majority (444, 72.5%) were increased in *E. lenta* cultures. Notably, the number of differentially abundant features at the final time point, both in total and among those assigned an identification, is substantially higher than previously reported metabolomic profiles of this species in ISP-2 and Mega media (Bisanz et al., 2020; Han et al., 2021) (**Figure S1C**). This increased sensitivity was expected given our use of both chemically defined culture media and untargeted metabolomics.

**Figure 1.**
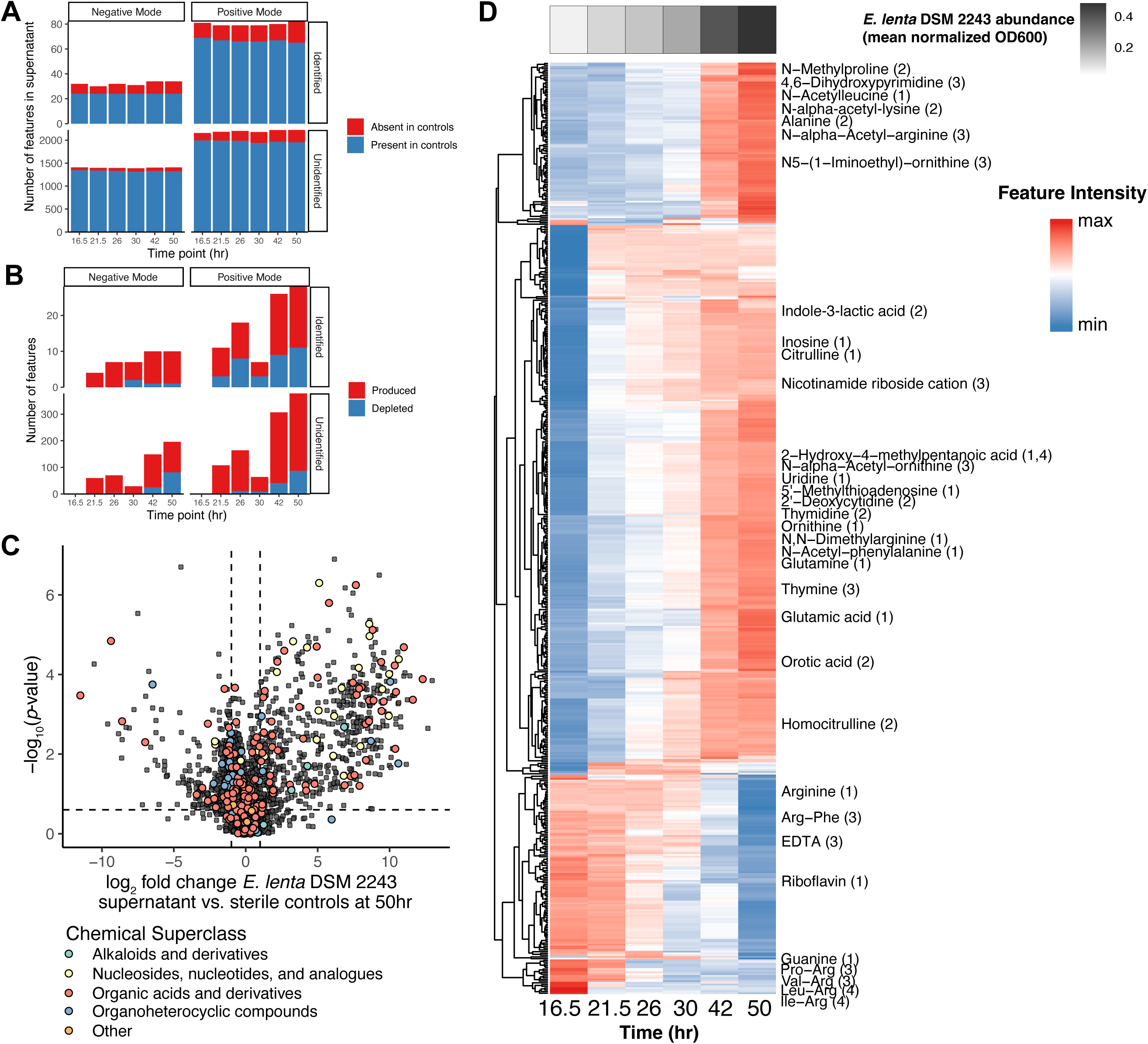
Production and depletion of diverse metabolites by *Eggerthella lenta* DSM 2243 in chemically defined media. **A)** Number of metabolite features detected by tandem LC-MS in culture samples at each time point. Features are considered present if their average peak height in supernatant is greater than 3x the average peak height in blank samples. Using both positive and negative ionization modes, an increasing number of features not found in controls appear in culture supernatants over time. **B)** Number of differentially abundant metabolite features compared with sterile control media at each time point, based on FDR-adjusted t-tests of log-transformed peak heights. **C)** Volcano plot of differentially abundant metabolite features at the final time point (50 hours) compared with sterile controls. *p*-values shown on the *y*-axis are based on Welch’s t-tests comparing values at the final time point vs. sterile controls (Benjamini-Hochberg adjusted). **D)** Heatmap of individual metabolite trajectories in cultures of *E. lenta* DSM 2243 grown in EDM1 batch culture. Features shown are those whose abundance was significantly different from controls (FDR-adjusted *p*<0.1 and absolute log_2_ fold change>0.75) at the final time point. Identified metabolites are labeled; the number in parentheses indicates the Metabolomics Standards Initiative confidence level for that identification (with 1 as highest confidence, see Methods). Values shown are average log-transformed peak heights, scaled for each feature. The gray heatmap at the top indicates the average batch culture density at each time point of *Eggerthella lenta* DSM 2243 in EDM1 (normalized OD600). See also **Figure S1-2, Table S1-2**.

Metabolites of diverse chemical classes are modified by *E. lenta* (Figure 1C-D). Compounds produced by *E. lenta* tended to be amino acid and nucleic acid metabolites. As expected, these included ornithine and citrulline, suggesting activity from the arginine deiminase pathway, which is highly expressed by *E. lenta* in the presence of arginine (Haiser et al., 2013). However, other arginine-related metabolites were also produced at lower levels, including N,N-dimethylarginine, N5-(1-iminoethyl)-ornithine, and homocitrulline, suggesting that arginine may also be metabolized via other pathways. Several other metabolites produced at lower levels appeared to be products of metabolism of other amino acids in the media, including 4-methyl-2-hydroxy-pentanoic acid (from leucine), indole-3-acetate and indole-3-lactic acid (from tryptophan), and 3-phenyllactic acid (from phenylalanine), consistent with one previous report of production of indole-containing compounds and phenyl acids by *E. lenta* (Beloborodov et al., 2009). Other metabolites produced in supernatants included the amino acids alanine, glutamate, glutamine, histidine, and lysine; as well as several intermediates in biosynthesis of both purines and pyrimidines (inosine, orotic acid, hypoxanthine, uridine, thymidine). Overall, the set of metabolites produced by *E. lenta* supports its previously reported dependence on arginine catabolism, but is highly multifaceted.

Of the 54 compounds in our EDM1 recipe, 22 were detected by untargeted metabolomics but just three were depleted significantly in *E. lenta* cultures (**Figure S1D**, Figure 1D): arginine, riboflavin, and EDTA (which is likely reduced due to complexing with metal ions rather than from direct uptake or metabolism). This result suggested that most compounds were included in excess, leading us to reduce the concentration of several non-depleted amino acids for subsequent experiments (**Table S1**). Interestingly, 5 of the identified metabolite features significantly depleted by *E. lenta* were not explicitly included in our defined media formulation, including guanine and five arginine dipeptides (Figure 1D). Since these compounds were found at low intensities, were annotated with high confidence, and are structurally related to intentionally included compounds, we inferred that they may be trace contaminants from commercial preparations of uracil and arginine (see *Methods*). Their rapid depletion indicates that their presence may influence growth and metabolic activity and reinforces the value of untargeted metabolomic profiling.

We examined the dynamics of metabolite production and depletion over the 50-hour growth of *E. lenta* in batch culture. Hierarchical clustering of metabolite trajectories indicated that among both produced and depleted features, some metabolites are produced/depleted rapidly early in growth while others shift more dramatically later as the culture approaches stationary phase (Figure 1D, **Figure S1E**). This observation suggests that two or more distinct growth phases may be occurring as resources are consumed from the media. Among identified metabolites, the trace guanine and arginine dipeptides are first depleted from the culture in early time points while citrulline, inosine, and indole-3-lactic acid are produced at relatively higher rates (Figure 1D). In the later phase, arginine is depleted more rapidly while alanine, 4,6-dihydroxypyrimidine, and various *N*-acetylated amino acid metabolites are produced.

To gain a better understanding of the contributions of individual nutrients to *E. lenta* growth, we systematically tested the effect of their removal from the media on growth of *E. lenta* DSM 2243 (*Methods*, **Table S2**). We collected growth curve data from EDM1 with and without each component and fit logistic growth models to the results, finding that 22 out of 41 compounds tested had a significant effect on at least one of the following growth parameters (Wilcoxon rank-sum test, FDR-adjusted *p*<0.2): carrying capacity (maximum density), growth rate, time to mid-exponential, and/or area under the growth curve (**Figure S2A**). The only compounds whose individual removal fully prevented growth of *E. lenta* were arginine, tryptophan, riboflavin, biotin, and magnesium (although it is plausible that other compounds are required in trace amounts and were not fully removed by our preparation methods, particularly minerals such as iron). In general, removing amino acids most commonly tended to reduce carrying capacity, consistent with a role as carbon and/or energy sources, while removing vitamins had more varied effects on the growth curve (**Figure S2B**).

### Acetate and arginine are key carbon and energy sources for *E. lenta*

Surprisingly, we found that sodium acetate contributed substantially to *E. lenta* growth in EDM1 (**Figure S2A**), even though it was included at a relatively low concentration (1 mM, compared to 57 mM arginine in EDM1). Since acetate is an abundant and variable metabolic byproduct of diverse human gut microbes (van der Hee and Wells, 2021), dependence on acetate could shape the ecological interactions of *E. lenta* in the human gut microbiota. Although our untargeted LC-MS workflow was not able to quantify acetate, we had observed accumulation of several *N*-acetylated compounds in supernatant (Figure 1D), suggesting that the amount of acetate incorporated into core metabolic pathways may be relatively small. However, acetate provided a dose-dependent increase in carrying capacity for *E. lenta* up to a concentration of at least 10 mM in EDM1 (Figure 2A). We therefore used a targeted derivatization and LC-MS/MS method to quantify acetate levels in supernatants from three strains of *E. lenta* (DSM 2243, AB8n2, and Valencia) grown in EDM1 with different acetate concentrations (0, 1, or 10 mM). Acetate was depleted to approximately the limit of quantification in cultures from the 1 mM acetate group, but not the 10 mM acetate group, confirming that a relatively small quantity is required for the observed level of *E. lenta* growth (**Figure S3A**). We tested the effect of replacing acetate with equimolar amounts of 10 other small carbon compounds, finding that no tested alternative compound provided a comparable benefit (Figure 2B). Based on these results, we chose to further investigate *E. lenta*’s acetate utilization pathways.

**Figure 2.**
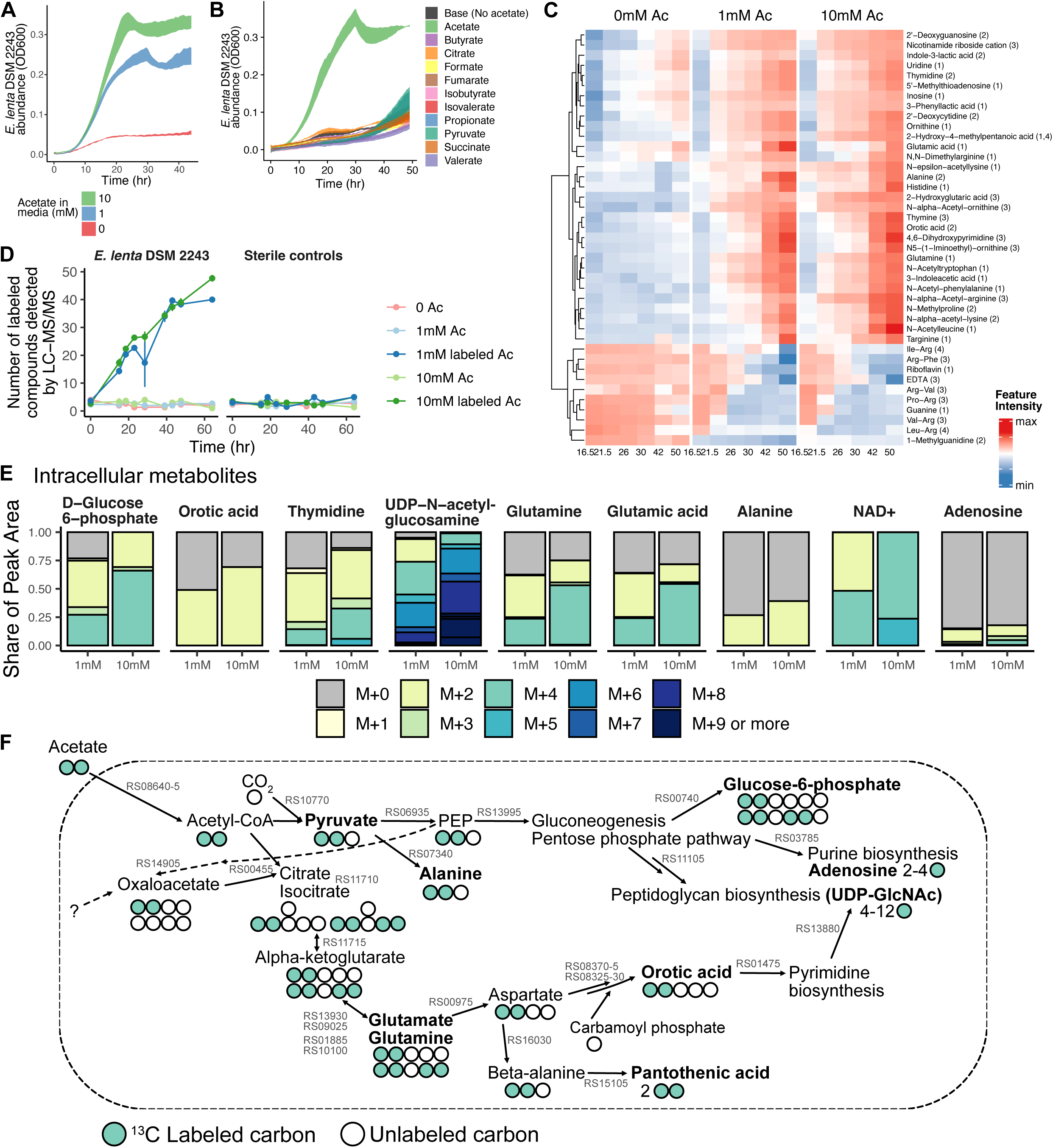
*E. lenta* uses acetate for nucleotide and peptidoglycan biosynthesis. **A)** Growth of *E. lenta* DSM 2243 in EDM1 media with varying concentrations of sodium acetate. **B)** Growth of *E. lenta* DSM 2243 in EDM1 media in which 1 mM sodium acetate is replaced with other small carbon compounds. **C)** Trajectories of identified metabolite features responsive to acetate concentration in *E. lenta* EDM1 batch cultures. Values are scaled average log-transformed peak heights from untargeted metabolomics profiling of supernatants. Labels show metabolite identity and MSI confidence level in parentheses. Metabolites shown are those that were assigned an identity and that had significantly different trajectories in the 0 mM vs. 1 mM acetate group based on spline regression comparison with the R package *santaR* (FDR-adjusted *p*<0.25). **D)** Stable isotope-resolved metabolomics profiling of *E. lenta* DSM 2243 in EDM1 media with ^13^C_2_ labeled acetate. The number of compounds with labeled isotopologues detected at a peak area > 10^5^ is shown for each sample group and time point, indicating incorporation of acetate into varied metabolites by *E. lenta*. **E)** Mass isotopologue distributions (MIDs) of intracellular metabolites. Each barplot shows the average isotopologue distribution in 1 mM and 10 mM acetate cultures. Compounds shown are those with an average labeled MID > 0.15 and a total peak area from labeled isotopologues of at least 10^4^ in at least one *E. lenta* DSM 2243 labeled acetate condition. **F)** Hypothesized pathways for incorporation of acetate into *E. lenta* central carbon metabolism and into biosynthetic pathways to produce labeled metabolites. Circles indicate the number of carbon atoms in selected compounds and are colored green to indicate incorporation of ^13^C isotopes from external acetate. Compound names in bold were detected with the observed labeling patterns in either intracellular metabolite extracts or culture supernatants. Corresponding enzymes are annotated in the *E. lenta* DSM 2243 genome for all reactions shown, and labeled in gray with the NCBI locus tag number. For pathways shown at a summary level (gluconeogenesis, pentose phosphate pathway, purine and pyrimidine biosynthesis, peptidoglycan biosynthesis), only the first enzyme in the pathway is labeled on the plot. See also **Figure S3-5**, **Data S1**.

First, we used our untargeted LC-MS metabolomics workflow to compare metabolites in supernatant over time from the same three *E. lenta* strains grown in EDM1 with different acetate concentrations (*E. lenta* DSM 2243 shown in Figure 2C, AB8n2 and Valencia in **Figure S3B-C**). Using smoothing spline models, we found that many produced or depleted compounds had significantly different abundance trajectories across the growth phase (FDR-adjusted *p*<0.25) depending on the presence of acetate. These included pyrimidine metabolites, *N*-acetylated amino acids, amino acid metabolites including indole-3-lactic acid and 2-hydroxyglutaric acid, and 423 unidentified metabolite features (Figure 2C). Of the 612 features produced by *E. lenta*, 53.4% had significantly different trajectories in the no acetate condition. Most differentially abundant compounds were associated with cell density and produced by *E. lenta* at higher levels when grown with higher acetate concentrations, reinforcing the general loss of biomass production in the absence of acetate.

To identify the specific pathways by which acetate is metabolized by *E. lenta*, we next profiled metabolites in the supernatant across time during growth of the same three strain isolates of *E. lenta* with ^13^C_2_ acetate provided as a stable isotope-labeled substrate (DSM 2243 in Figures 2D-F, 2 additional strains in **Figure S4**). We detected the incorporation of ^13^C labeled atoms in 52 features in *E. lenta* supernatants at the final time point, of which 24 were previously identified as responsive to acetate concentrations (*Methods,* Figures 2D, **S4**). Acetate was incorporated into diverse products across metabolite classes, but was found at the highest enrichment levels in nucleotide and carbohydrate metabolites (**Data S1**).

Because many core metabolites are not produced in excess or secreted during growth, we also analyzed intracellular metabolites from extracts collected at a single time point in the late-exponential growth phase. Labeled intracellular compounds included glutamate, glutamine, sugars, nucleotide metabolites, and UDP-*N*-acetyl-glucosamine, a primary component of peptidoglycan (Figure 2E), as well as seven labeled compounds of unknown identity. The signal from carbohydrate-related compounds including glucose-6-phosphate and UDP-*N*-acetyl-glucosamine was almost exclusively from labeled isotopologues (97.5% in 1 mM acetate and 100% in 10 mM acetate), indicating that synthesis of these compounds using acetate may be more efficient than any alternative non-acetate-dependent pathways available to *E. lenta* in the EDM1 condition.

Acetate-derived extracellular and intracellular metabolites were consistent across the two additional strains of *E. lenta*. While the overall rate of acetate incorporation differed between the three strains, the set of extracellular and intracellular labeled compounds was fully consistent. Isotopic enrichment for two additional extracellular metabolites (malonic acid and 3-hydroxy-myristic acid) was identified in both of these strains as well as four additional intracellular metabolites in one or both strains (all of unknown identity), confirming that acetate is incorporated by *E. lenta* into varied biosynthetic pathways (**Figure S4**).

Based on these results and metabolic gene annotations of the *E. lenta* DSM 2243 genome, we hypothesized that *E. lenta* converts acetate to acetyl-CoA via acetate kinase (ELEN_RS08645) and phosphate acetyltransferase (ELEN_RS08640). Acetyl-CoA could then be used as a carbon source via two routes: conversion to glutamate by a partial citric acid cycle, and synthesis of pyruvate by the enzyme pyruvate-ferredoxin oxidoreductase (PFOR, ELEN_RS10770) (Figure 2F). This hypothesis is consistent with the organization of the *E. lenta* DSM 2243 genome, as two of the three enzymes required for conversion of acetyl-CoA to glutamate are co-located (aconitate hydratase and isocitrate dehydrogenase, ELEN_RS11710, ELEN_RS11715). Genes for another partial component of the citric acid cycle—fumarate hydratase and malate dehydrogenase—are co-located in another region of the genome (ELEN_RS056[70-90]), suggesting they may act in a separate functional role. Taken together, these data suggest that *E. lenta* uses acetate as a key carbon source for synthesis of biomass components, in tandem with ATP generation from arginine catabolism, anaerobic respiration, and/or other unknown pathways.

However, we inferred that acetate is likely not the sole carbon source used by *E. lenta* in EDM1, given the relatively low concentration required for growth promotion and the abundance of unlabeled isotopologues detected for many produced compounds (**Data S1**). We wondered whether arginine or ornithine may also be substrates for synthesis of biomass components, or if arginine is exclusively catabolized to ornithine for ATP production, as suggested by one previous study in rich media (Sperry and Wilkins, 1976). We first confirmed that citrulline, but not ornithine, can replace arginine with nearly equivalent growth in EDM1, replicating a previous result in rich media [(Haiser et al., 2013), **Figure S5A**]. We then analyzed intracellular and extracellular metabolites from *E. lenta* DSM 2243 growing in EDM1, this time with ^13^C_6_ L-arginine as a stable isotope-labeled substrate. We found by far the largest composition of ^13^C enriched isotopologues in ornithine, citrulline, and other closely related compounds (**Figure S5B-E**), indicating that arginine is predominately processed by the arginine deiminase pathway. However, we observed M+1 enrichment (i.e. incorporation of a single ^13^C carbon atom from arginine) in produced glutamine, orotic acid, and pyrimidines, among others (**Figure S5C-D**), suggesting biosynthesis from the carbamoyl phosphate intermediate. Labeled M+5 isotopologues of proline and prolinamide also appeared at low levels at later time points, likely indicating a slower flux producing these compounds from accumulated ornithine (**Figure S5D-E**). Yet in total, only 29/324 features were detected with ^13^C enrichment for five or more carbon atoms in intracellular extracts, and most appeared closely related to arginine, citrulline, and ornithine (**Data S1**). These results confirm that arginine is primarily an energy source and not a major biosynthetic precursor for *E. lenta* (**Figure S5F**).

### A genome-scale metabolic model of the *E. lenta* type strain recapitulates growth, metabolite, and gene expression phenotypes

COnstraint-based Reconstruction and Analysis (COBRA) is a set of computational tools that has been applied to interpret -omics data and optimize metabolic activities for various microbes of importance in basic science, metabolic engineering, and medicine (Gu et al., 2019; Monk et al., 2017; Zhang et al., 2017). It has been proposed as a promising strategy to predict phenotypes and design modifications to complex host-associated microbial communities by synthesizing information about the physiology of individual members and the available nutrients into a rational framework (Chiu et al., 2014; Diener et al., 2020; Hertel et al., 2019). However, the value of such a framework is dependent on its ability to accurately describe the contributions of metabolically active community members. The reconstructions currently available for many anaerobic microbes have only been curated to a limited degree and remain minimally validated. Therefore, we used our *in vitro* platform to curate and analyze a genome-scale metabolic network model of *E. lenta* DSM 2243 growth in EDM1 and assessed the degree to which this model can explain *E. lenta* metabolic phenotypes across conditions.

We obtained a genome-scale metabolic reconstruction from the AGORA database version 2.0.0 (Heinken et al., 2020), which we term iEL2243_2. Initial testing indicated that the model was incapable of biomass production in EDM1 media, so we performed additional curation of model reactions and transporters (**Table S3**). We curated the reconstruction based on genome annotations from multiple sources (Henry et al., 2010; Pascal Andreu et al., 2021; Price et al., 2022) and added transporters for strongly depleted and produced compounds that were identified with high confidence in our metabolomics data. Throughout this process, we compared model results with experimentally observed growth in chemically defined media conditions, using these results to inform the curation process and add missing reactions where supported by experimental data. We simulated metabolic fluxes in different conditions by converting media concentrations into estimated maximum nutrient uptake rates for each compound. While these models are typically validated by comparison with gene essentiality data (Thiele and Palsson, 2010), the tools to generate such data are not yet available for *E. lenta.* We instead evaluated whether the model was consistent with observed metabolite utilization and production and with gene expression during exponential growth in EDM1, and whether predicted essential genes were conserved across strain genomes.

This process resulted in a model with 1,244 reactions linked to 727 gene annotations and 1,218 metabolites (Figure 3A). The largest number of reactions were in the subsystems of fatty acid synthesis, extracellular transport, and glycerophospholipid biosynthesis (Figure 3B). Flux balance analysis of the final model estimated the maximum growth rate of *E. lenta* DSM 2243 in EDM1 to be 0.96 hr^-1^, higher than experimental values (median 0.32 hr^-1^, **Figure S2B**). The existence of a difference between these values is not surprising given that organisms do not necessarily grow at their theoretical maximum growth rate, and growth constraints may exist that are not encoded in the metabolic network model (Thiele and Palsson, 2010). However, the relatively large discrepancy indicates that additional modifications to the biomass equation may further improve the model.

**Figure 3.**
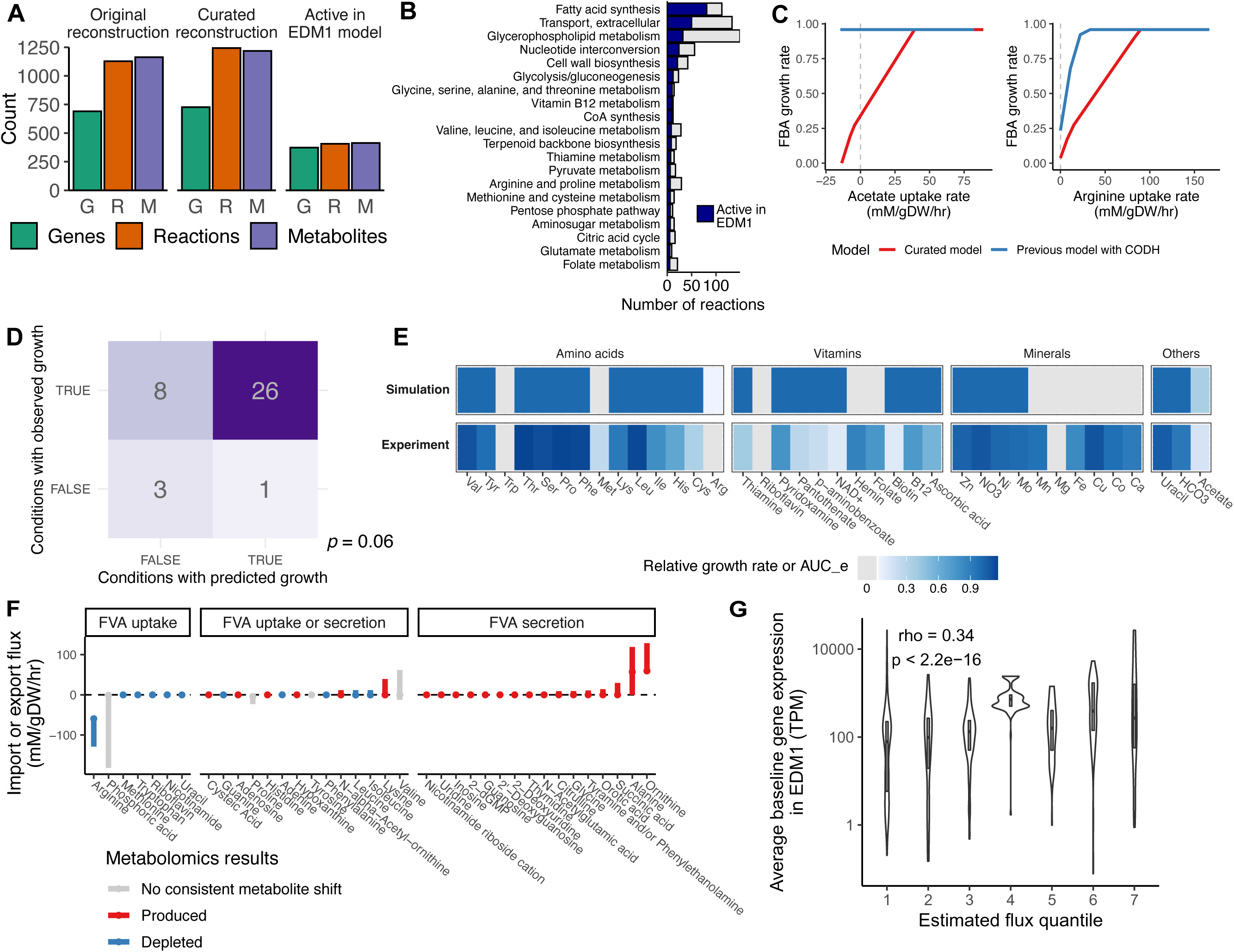
A curated genome-scale metabolic model of *E. lenta* DSM 2243 partly explains growth phenotypes across conditions. **A)** Summary of the curated reconstruction of *E. lenta* DSM 2243 indicating the number of genes, reactions, and metabolites in the original and curated models, and the share of those required to be active for growth in EDM1 based on parsimonious flux balance analysis (pFBA). **B)** Summary of the total number of reactions by subsystem, and the share of each subsystem predicted to be active in EDM1 (only the top 20 subsystems are shown). **C)** Acetate and L-arginine uptake dependencies inferred by the model. In the final curated model (red lines), the maximum growth rate decreases with decreasing availability of both L-arginine and acetate, qualitatively consistent with experimental data. A previous model incorporating a carbon monoxide dehydrogenase reaction based on (Harris et al, 2018) (blue lines) failed to recapitulate the expected dependencies. **D)** Confusion matrix summarizing a comparison of growth/no growth between the iEL2243_2 model vs. experimental observations for leave-one-out media conditions. **E)** Full set of quantitative comparisons underlying panel D. Each column shows the FBA-inferred maximum growth rate in the EDM1 condition with a media component removed, paired with the experimentally observed area under the empirical growth curve for that condition. A gray tile indicates zero growth. **F)** Comparison of shifts in metabolomics data with uptake and secretion rate ranges inferred for the same compounds by flux variability analysis (FVA). Metabolites that can only be imported according to FVA were decreased in metabolomics data, while those with potential for being produced were indeed produced. **G)** Comparison of absolute fluxes inferred by pFBA with gene expression of linked enzymes of *E. lenta* DSM 2243 during exponential growth in EDM1. Within flux quantiles (on the x-axis), genes are expressed at a wide range of levels, but genes linked to reactions with the highest fluxes are generally highly expressed. See also **Figure S6**, **Table S3-4.**

The initial model with nonzero growth in EDM1 did not recapitulate the experimentally observed dependencies on either arginine or acetate (Figure 3C). We noticed that this lack of dependency was linked to the inclusion of Wood-Ljungdahl acetogenesis reactions in the model, previously suggested to be present in *E. lenta* (Harris et al., 2018; Hylemon et al., 2018). The presence of these reactions allowed the model to draw on an effectively unlimited source of acetyl-CoA from CO_2_ and H_2_. Regardless of whether the previous annotation of this pathway (which has not been biochemically validated) is correct, reductive acetogenesis may not be thermodynamically favorable during *in vitro* growth in our anaerobic chamber, where the H_2_ concentration is ≤ 5% (Smith et al., 2020). Blocking model flux through the carbon monoxide dehydrogenase reaction of this pathway increased growth dependency on uptake of both arginine and acetate, reflecting our experimental observations (Figure 3C). The model also found no growth benefit from pyruvate, citrate, and other fatty acids based on a lack of annotated transporters for these compounds, consistent with experimental results.

In another key curation step, required to enable biomass production by the model in EDM1, we noticed that *E. lenta* lacks an annotated gene for the enzyme enoyl-acyl protein carrier reductase, which performs the elongation in the typical type 2 fatty acid synthesis pathway used in bacteria. Because fatty acid biosynthesis is essential and previous studies have noted a high level of diversity in this essential step among bacterial genomes (Massengo-Tiassé and Cronan, 2009), we preserved this step in the model without any current gene annotation. This gap may indicate a novel enzyme family performing this conversion (**Table S3**).

We applied the iEL2243_2 model to predict growth phenotypes across our leave-one-out chemically defined media conditions, finding that these were generally consistent with some remaining notable exceptions (Figure 3D, overall Matthews correlation of 0.35, Fisher exact test odds ratio=9.1, *p=*0.06). Amino acid dependencies matched well between the model and experimental data, with the exception of cysteine, which likely provides a benefit as a reducing agent that is not accounted for by the model (Strobel, 2009). Vitamin dependencies were also generally consistent, with the notable exception of folate, which had no effect on growth despite the lack of several genes for reactions in the canonical folate biosynthesis pathway and the absence of a known dihydrofolate reductase enzyme (Rodionov et al., 2019). The phenomenon of presumed-essential but absent folate genes in bacterial genomes has been recognized previously (de Crécy-Lagard et al., 2007; Levin et al., 2004; Rodionov et al., 2019), suggesting the possible existence of undiscovered alternative enzymes. Notably, growth was negatively affected by the removal of the folate precursor *p-*aminobenzoate (**Figure S2A**). Most of the remaining discrepancies between the model and the growth data are in conditions in which metal ions were removed, which were expected to be required by the model (Cu^2+^, Ca^2+^) but were not essential based on our experiments (Figure 3E). However, these likely reflect difficulties in fully removing trace minerals in our experiment rather than errors in the model reconstruction.

While we curated the model based on growth data, we did not incorporate our metabolomics data except to add transporters for highly differentially abundant metabolites. Even so, we found that there was a high correspondence between observed metabolite shifts and the possible uptake and secretion fluxes inferred by flux variability analysis (FVA) of the model. FVA identifies the range of fluxes for each reaction that are compatible with near-maximum growth. All 37 identified metabolites present in both the model and our metabolomics data displayed experimental shifts in abundance qualitatively compatible with inferred flux ranges (Figure 3F), providing additional support for model quality.

We further compared the iEL2243_2 inferred flux profile with RNA-Seq data from *E. lenta* growing in this condition, which was not used for model curation (*Methods*). 71.9% of genes linked to active reactions were in the top half of metabolic genes by expression level in the EDM1 condition (> 109 transcripts per million), and 91.9% were in the top 75%. Expression level and absolute flux magnitude were highly correlated across all genes linked to metabolic reactions (Spearman rho=0.34, *p*<2.2×10^-16^, Figure 3G). While we would not expect a perfect correlation between expression and metabolic flux, correspondence between the two provides support that our model has correctly identified pathways with high activity.

Having established consistency with experimental data, we next examined overall reaction fluxes and key pathways in the final model. We found that fewer than half of reactions were predicted to be active in EDM1 by parsimonious flux balance analysis (pFBA, Figure 3A). In the pFBA solution, acetate is incorporated into a partial reductive citric acid cycle via pyruvate formate oxidoreductase (PFOR), which then feeds lipid and carbohydrate biosynthesis pathways, consistent with our SIRM results and with our RNA-Seq data, where PFOR was one of the most highly expressed genes. The vast majority (99.6%) of arginine uptake flux was directed to ATP generation, and 58.4% of ATP generation was sourced from the arginine deiminase pathway (which contains the 1st, 3rd, 4th, and 5th most highly expressed protein-coding genes in our RNA-Seq data, **Table S4**). The remainder of ATP generation in the pFBA solution was attributed to anaerobic respiration via an ATP synthase reaction, although the specific electron transport chain substrates were not clear. However, consistent with this hypothesis, genes linked to respiration were expressed at moderate levels, including ATP synthase subunits and an Rnf electron transport complex, and *E. lenta* is known to have a large number of poorly characterized enzymes potentially involved in electron transfer (Maini Rekdal et al., 2020; Ravcheev and Thiele, 2014). The model also identified the regeneration of NADP+ via transaminase reactions (using mainly pyruvate and/or branched chain amino acids) and glutamate dehydrogenase as a key high-flux pathway.

Finally, we applied the model to predict the effects of knocking out individual reactions on growth of *E. lenta*. 15.3% of all reactions in iEL2243_2 were predicted to be essential in any condition and 19.4% to be essential in EDM1. These reactions tended to be involved in lipid metabolism, cell wall biosynthesis, and transport of essential metabolites (**Figure S6A**). Genes linked to reactions whose removal reduced growth to < 70% of wild type levels were found in a greater number of *E. lenta* strain genomes than other genes (Wilcoxon rank-sum test, *p*=0.001, **Figure S6B**) and were more likely to be part of the core genome (found in all strains; Fisher exact test odds ratio = 1.74, *p*=0.0002). Overall, while significant manual curation was required for the model to recapitulate realistic growth in EDM1, our updated model is able to predict and interpret many aspects of *E. lenta* growth and metabolic activity across conditions.

### The strain-variable *E. lenta* metabolome is enriched for nucleotides and cell wall metabolites and can be linked to genome variation

Our initial efforts to characterize *E. lenta* core metabolism focused mainly on the type strain. However, *E. lenta* has an open pan-genome and established variability in secondary and xenobiotic metabolism (Bisanz et al., 2020). We therefore evaluated the extent to which the metabolic profile of this species is conserved across a larger number of strain isolates. We used untargeted metabolomics to profile stationary phase supernatants of 30 strains grown in EDM1 (**Figure S7A-B**) and used linear models to identify features with significant strain-associated differences in abundance. Over half of the features produced by the UCSF DSM 2243 type strain (52.8%) were variable across strains of *E. lenta* (Figure 4A), and 1,097 features produced by at least two other strains were not produced by the type strain. Divergence in metabolite profiles between strains was not associated with phylogenetic divergence based on an alignment of core genes (Procrustes analysis, *p*=0.31, Figure 4B), consistent with previous findings from untargeted metabolomics profiling of these strains in rich media with a different metabolomics platform (Bisanz et al., 2020). Overall metabolite profiles were moderately associated with presence/absence patterns of variable gene families between strains (*p=*0.03, Figure 4B), indicating that the presence or absence of biosynthetic genes and pathways only partly explains variation in the metabolome and that other factors like gene regulation and enzymatic activity may also play a substantial role.

**Figure 4.**
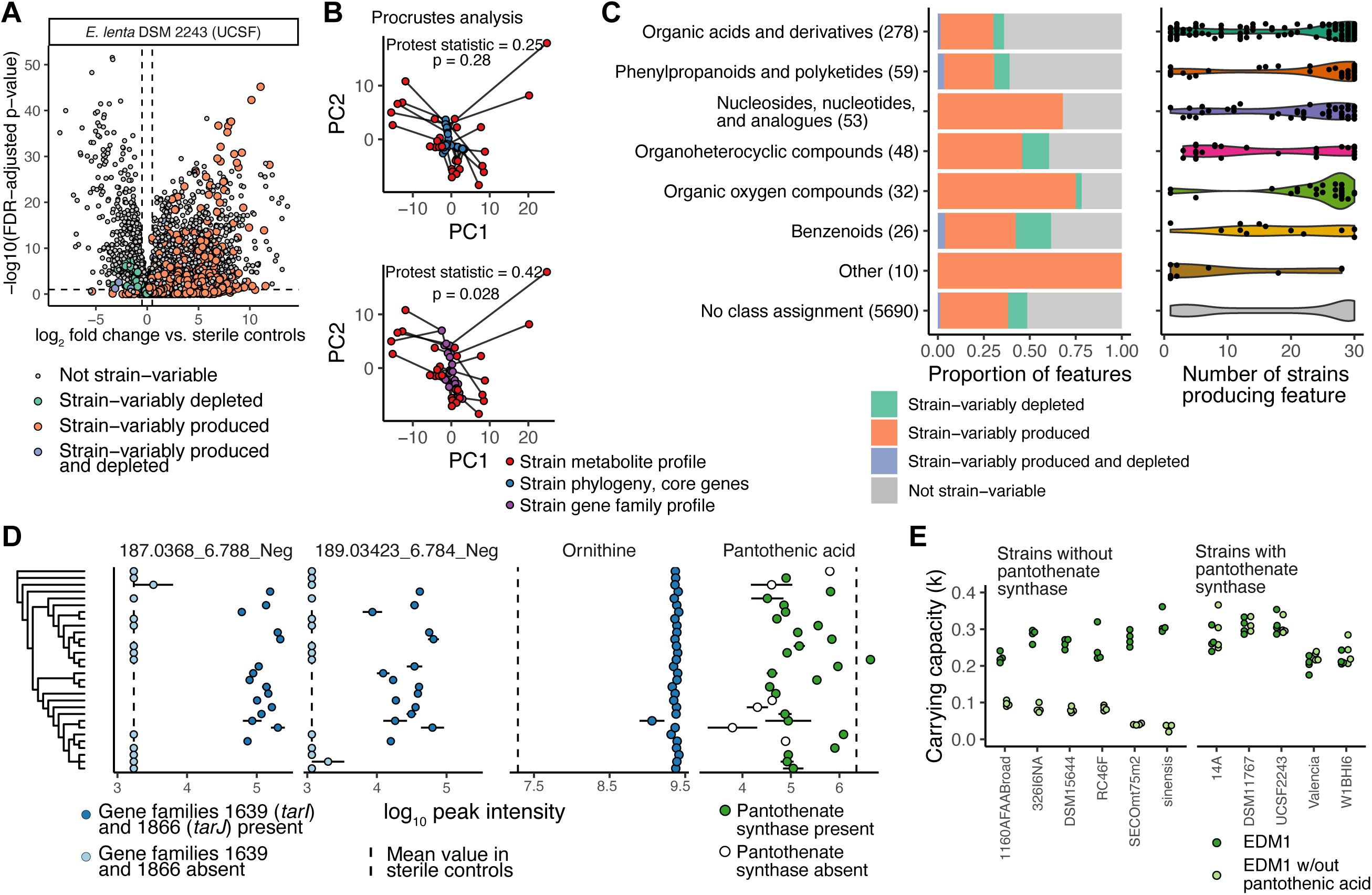
Extensive within-species variation in *E. lenta* metabolites can be linked to variable gene families. **A)** Volcano plot of metabolite features detected in stationary phase supernatants of *E. lenta* DSM 2243 (UCSF lab strain) vs. sterile controls. *P-* values are based on Benjamini-Hochberg corrected Welch’s t-tests. Features are colored based on whether their classification as significantly produced or depleted (increased or decreased with FDR-adjusted *p*-value<0.1 and log2FC>0.5) is consistent across 28 other *E. lenta* isolates and one isolate of *Eggerthella sinensis* profiled in the same experiment. B) Procrustes analysis of overall metabolite profiles compared with genome features. The upper plot shows a rotated Procrustes superimposition of average metabolite profiles for each isolate (red points) and the phylogenetic distance between them based on an alignment of core genes (blue points). The lower plot shows a superimposition of metabolite profiles and profiles based on the presence/absence of variable gene clusters (purple points). C) The left-hand panel shows the distribution of strain-variable features in various ClassyFire chemical superclasses, based on Feature-based Molecular Networking with GNPS. The number in parentheses for each class indicates the total number of features with that assignment. The right-hand panel shows the number of strains producing a given feature, among features produced by any *Eggerthella* isolate. Each point represents a single feature, and its position on the x-axis indicates the number of strains for which that feature was significantly increased (FDR-adjusted *p*-value < 0.1 and log_2_ fold change > 0.5) in supernatants compared with controls. Superclasses (y-axis labels) are the same as in panel C. E) Feature abundances of example metabolites across strains. The first two panels show two strain-variable unidentified features associated with the presence of specific strain-variable gene families - putatively identified as the two dominant naturally occurring isotopes of an [M+Cl-] adduct of the teichoic acid component ribitol. The points indicate the log-transformed abundances of these features for each strain. The dotted line in each panel indicates the average level of that feature in sterile controls. Points in dark blue represent strains whose genomes contain genes for a ribitol-5-phosphate cytidylyltransferase (*tarJ*) and ribulose-5-phosphate reductase (*tarI*) not found in other genomes. The third and fourth panels show a highly conserved identified metabolite (ornithine) compared with a strain-variable identified metabolite (pantothenic acid). Points in white in the pantothenic acid panel indicate strains whose genome lacks the final step in the biosynthetic pathway for this metabolite. Points are shown as mean and standard error across three replicates. The order of strains on the y-axis matches their phylogeny, shown in Figure S7A. F) Strains lacking a gene annotated as pantothenate synthetase deplete pantothenic acid completely from media (previous panel) and have a substantial growth defect when grown in the absence of pantothenic acid (left panel). Closely related strains that do possess this gene are unaffected by removal of pantothenic acid (right panel). Carrying capacity is estimated based on a logistic growth model fit by the R package *growthcurver.* See also Figure S7, Table S5-6.

While strain-variable metabolites were quite diverse, they were enriched for certain chemical classes. 92.0% of strain-variable metabolites had no identity information, a similar ratio to the total number of metabolite features (91.8% of features in the whole dataset). Among other features, organic acids (which included many amino acid metabolites) were the least likely to be strain-variably produced. In contrast, organic oxygen compounds (which included several features identified as sugars) and nucleotide metabolites were more likely to be strain-variably produced, and organic heterocyclic compounds and benzenoids were enriched for strain-variable depletion (Figure 4C). The share of strains producing any individual feature varied widely (Figure 4C), although the largest number of features (76.8%) were produced by either only a few (<4) strains or nearly all (>27) strains (**Figure S7C**).

Given the large share of unidentified metabolites in our dataset, we evaluated whether linking strain-variable metabolites with strain-variable genes could inform metabolite annotations. We performed an association analysis between metabolite feature abundances and the presence of specific accessory gene families, applying a method developed for previous analysis of this *E. lenta* strain collection (Bisanz et al., 2020). A full 39.0% of metabolite features were significantly associated with the presence of one or more variable gene families (FDR-adjusted *p*<10^-4^). Using stricter filtering criteria for significance, effect size, and separability, 84 metabolite features (1.3%), of which 80 had no annotation, were linked with the presence of variable genes (**Table S5**, *Methods*). Gene families linked to these features were enriched for KEGG annotations in sulfur metabolism (*q* = 0.00017), ABC transporters (*q* = 0.02), porphyrin metabolism (*q* = 0.03), and biosynthesis of nucleotide sugars (*q* = 0.049), consistent with the profile of identified variable metabolites.

As a case study, we further examined two of the top hits from this analysis, two closely related but unidentified metabolite features highly associated with the presence of two adjacent gene families (Figure 4D). These gene families were annotated by Prokka (Seemann, 2014) as ribulose-5-phosphate reductase 1 (*tarI*) and a ribitol-5-phosphate cytidylyltransferase (*tarJ*), which are essential enzymes in the biosynthesis of CDP-ribitol teichoic acid. Teichoic acids are an abundant component of the cell wall of gram-positive bacteria that can take multiple forms and can be synthesized with either CDP-glycerol or CDP-ribitol subunits (Brown et al., 2013; Percy and Gründling, 2014; Weidenmaier and Peschel, 2008). Interestingly, the *m/z* value and MS2 spectrum of the linked features were consistent with an annotation as the two dominant [M+Cl]-naturally occurring isotope adducts of a 5-carbon sugar alcohol - *i.e.* potentially ribitol, xylitol, or a related compound.

Further examination of the *tar/tag* biosynthetic gene cluster in which these genes are located revealed extensive strain diversity, with 10 different gene arrangements across the 30 isolates (**Figure S7D**), suggesting recent positive selection possibly as a form of phage defense (Buttimer et al., 2022; Soto-Perez et al., 2019) or host immune interaction (van Dalen et al., 2020). Most genomes have one or more genes with homology to *E. coli arnC* genes in this region, indicating that the products may be lipoteichoic acids anchored to the cell membrane rather than wall teichoic acids (Percy and Gründling, 2014). Among *E. lenta* genomes without *tarI* and *tarJ*, all except the type strain have a *tagD* gene in the same region instead, which catalyzes the synthesis of CDP-glycerol subunits instead of CDP-ribitol (**Figure S7D**) and would be consistent with the absence of extracellular ribitol in those strains. Two other metabolite features were associated with the presence of other members of this gene cluster, possibly indicative of other strain-variable cell wall components (**Table S5**). This example illustrates that comparative multi-omics can be a powerful strategy to identify and begin to decipher the functional consequences of strain variation, even when metabolite identities are not confirmed.

In addition to the unbiased association analysis above, we also assessed whether strain variation in metabolites of known identity could be predicted based on relevant gene annotations. We created genome-scale metabolic reconstructions of a subset of strains included in this experiment (n = 24, using the DEMETER pipeline), curated them using a limited version of the process applied to the type strain (*Methods*), and again predicted growth and reaction fluxes in EDM1 and in leave-one-out media conditions using flux balance analysis. Across the metabolic networks of *E. lenta* strains, most reactions were conserved, including arginine metabolism and central carbon metabolism (**Table S6**). Tryptophan and riboflavin auxotrophies were also predicted to be conserved across strains. Variable reactions tended to be in the subsystems of transport, fatty acid biosynthesis, cell wall biosynthesis, and nucleotide interconversion (**Figure S7E**). Consistent with the central role of arginine metabolism, ornithine and citrulline levels in our metabolomics dataset were very consistent across strains. Ornithine was among the least variable metabolite features (Figure 4D), and one of the most correlated with biomass (as estimated by optical density, Spearman rho=0.36, FDR-adjusted *p=*0.1).

While the predicted effects of most compounds on growth were similar or identical across strains (**Figure S7F**), we noticed a clear difference in pantothenic acid dependence, as a subset of strains were predicted to be unable to grow in its absence. These strains lack the final enzyme in the biosynthesis pathway for pantothenic acid, which is itself a precursor of coenzyme A. Pantothenic acid was depleted to varying degrees in our metabolomics data, reaching the lowest levels in strains that lack pantothenic acid synthase (Figure 4D). Notably, M+2 isotopologues of pantothenic acid were also detected in supernatants from the acetate SIRM experiment, corroborating that at least three *E. lenta* strains synthesize this vitamin *de novo* (**Data S1**, **Figure S4B,** Figure 2F). We tested growth of pantothenate synthase-lacking strains in comparison with a subset of genetically similar strains in EDM1 with or without pantothenic acid, confirming that strains without this gene family had a greatly reduced carrying capacity in the absence of pantothenic acid (Figure 4E) and highlighting the ability of curated genome-scale models to predict phenotypic differences. Overall, our analysis of strain variation in metabolite profiles is consistent with a model in which *E. lenta*’s distinctive central carbon and energy metabolism is a core species trait, while more peripheral biosynthetic pathways including synthesis of cofactors and cell surface components can vary freely to adapt to specific microenvironments (Monk et al., 2013).

### Comparison of *E. lenta*’s metabolic profile *in vitro* and *in vivo* identifies shared signatures and usage of a novel nutrient

Having characterized the metabolic profile of the *E. lenta* species in a simplified *in vitro* environment, we next asked how these findings compare with its metabolic activity in a host, and whether our *in vitro* platform could help identify metabolic processes performed by *E. lenta* within the gastrointestinal tract. We monocolonized germ-free (GF) mice with one of three strains of *E. lenta* by oral gavage, collected serum and intestinal contents after two weeks of colonization, and profiled metabolites using the same LC-MS/MS untargeted metabolomics workflow as above. We identified features that were significantly differentially abundant in *E. lenta*-colonized mice vs. their GF counterparts using linear mixed models and compared these features with our *in vitro* metabolomics datasets, identifying metabolites consistently shifted by the presence of *E. lenta* across environments. After data processing, quality filtering and dereplication, we obtained a dataset of 19,714 metabolite features from intestinal samples. Of these, 16.7% were significantly differentially abundant (FDR-adjusted *p*<0.2) in response to colonization with at least one strain in at least one segment of the intestinal tract, indicating a substantial metabolic impact of *E. lenta* on the intestinal environment (**Figure S8A**). Interestingly, despite previous data showing colonization of *E. lenta* DSM 2243 at similar levels from the ileum to the colon in GF mice (Bisanz et al., 2020), only 1.6% of features were significantly shifted in the ileum, compared with 11.5% in colon and 8.1% in the cecum. Additionally, only 21 features (0.41%) were differentially abundant in serum in response to any of the three strains. Overall separability of metabolite profiles between germ-free and colonized was also highest in the cecum and colon (Figure 5A). These results indicate that *E. lenta*’s strongest metabolic effects are restricted to the lower intestinal tract.

**Figure 5.**
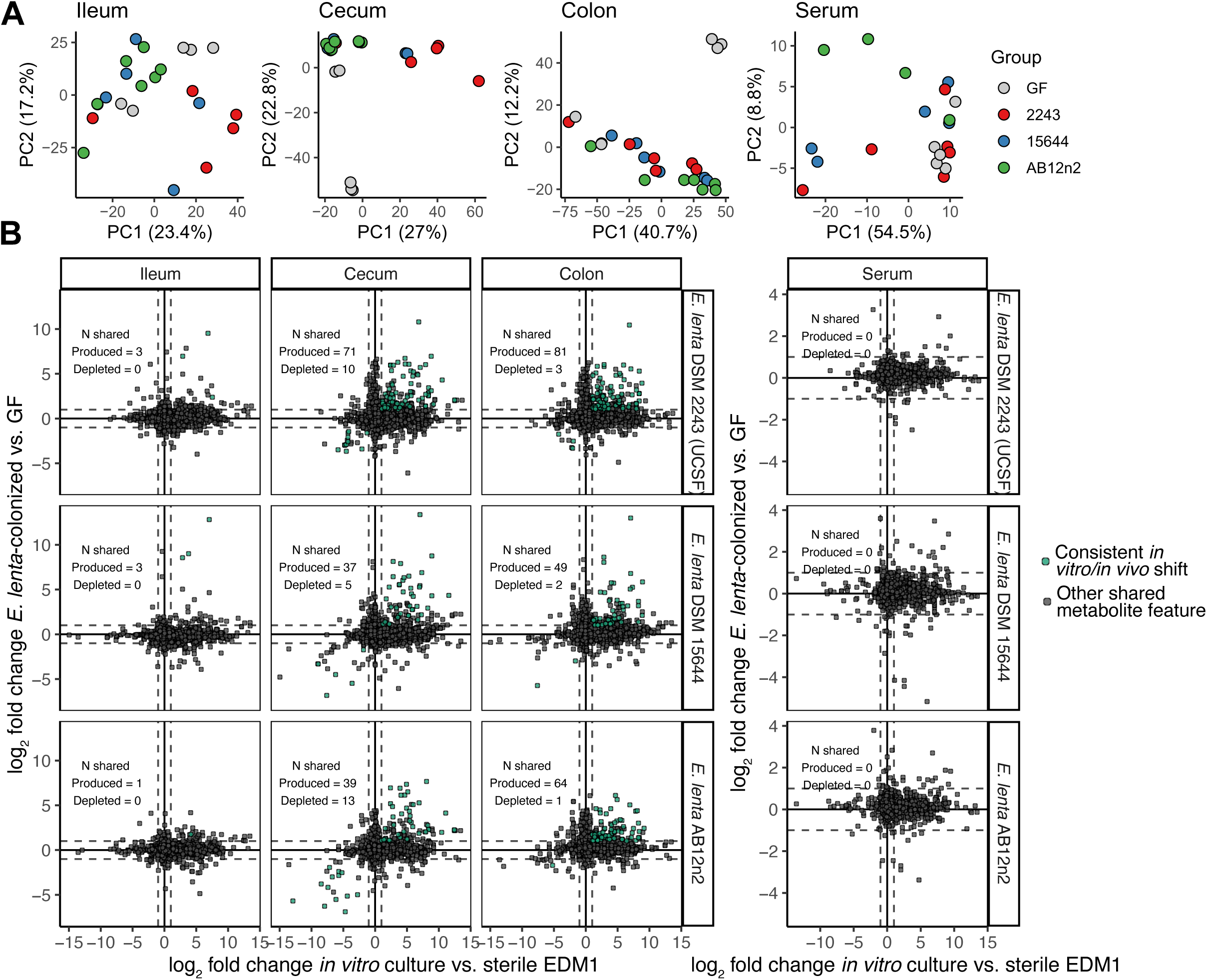
Comparison between *E. lenta*’s metabolic footprint *in vivo* and *in vitro* reveals shared metabolite signatures. **A)** Principal component analysis of untargeted metabolomics profiles of intestinal contents and of serum. **B)** Comparison of the effect of *E. lenta* on metabolite features detected in both EDM1 cultures and monocolonized mice. Each point represents a metabolite feature detected in both datasets. The *x-*axis indicates the log_2_ fold change of each feature in supernatants compared with sterile controls, compared with the estimated log_2_ fold change of that feature in monocolonized mice compared with germ-free mice. Points are colored green if the feature is significantly differentially abundant in gnotobiotic mice and is shifted in the same direction by the corresponding strain in the stationary phase *in vitro* experiment. See also **Figure S8**, **Data S2.**

We assessed the extent to which metabolite features produced by *E. lenta* in cell culture are detectably shifted by the presence of *E. lenta* in mice. To do so, we integrated our processed metabolomics datasets by linking metabolite features across datasets with highly similar *m/z*, RT, and MS2 spectra (see *Methods*). Based on this analysis, 37.2% of identified metabolite features in intestinal contents and 12.2% of features overall were also detected *in vitro* (**Figure S8B**). We compared the estimated log_2_ fold change of each linked feature *in vitro* with the corresponding shifts *in vivo* (full set in **Data S2**; Figure 5B shows the comparison with the strain collection dataset in Figure 4, **Figure S8C** shows a comparison with the dataset in Figure 1). 202 features significantly increased by the presence of *E. lenta* DSM 2243 in cecal contents were also increased in one of our EDM1 *in vitro* datasets, providing support that they are directly produced by *E. lenta in vivo*. These features represented 78.9% of the set that could be linked across datasets and 20.9% of the full set of *E. lenta* DSM 2243-increased features in cecal contents. Only 18 metabolites depleted in cecal contents were similarly depleted *in vitro*, but only three of the other 405 depleted features were detected *in vitro* at all, indicating that *E. lenta* likely uses a much richer set of nutrients *in vivo* than those available in EDM1. Overlapping produced and depleted metabolites were found in greater abundance in the cecum and colon than the ileum and serum (Figure 5B), again suggesting a greater metabolic footprint of *E. lenta* in the lower gastrointestinal tract relative to other sites.

Ornithine was among the most increased features across sampling sites and strains, consistent with our *in vitro* data (Figure 6A, **Figure S9A**). Other consistently increased features included 5-methyluridine, citrulline, glutamine, and lysine (**Data S2**). Interestingly, arginine was only significantly reduced by colonization with one of the three *E. lenta* strains in this experiment (**Figure S9B**). However, most other proteinogenic amino acids were increased in abundance in intestinal contents in colonized mice compared with GF (**Figure S9C**), likely due to differences in host activity, so the absence of an increase in arginine may be consistent with arginine usage by *E. lenta*.

**Figure 6.**
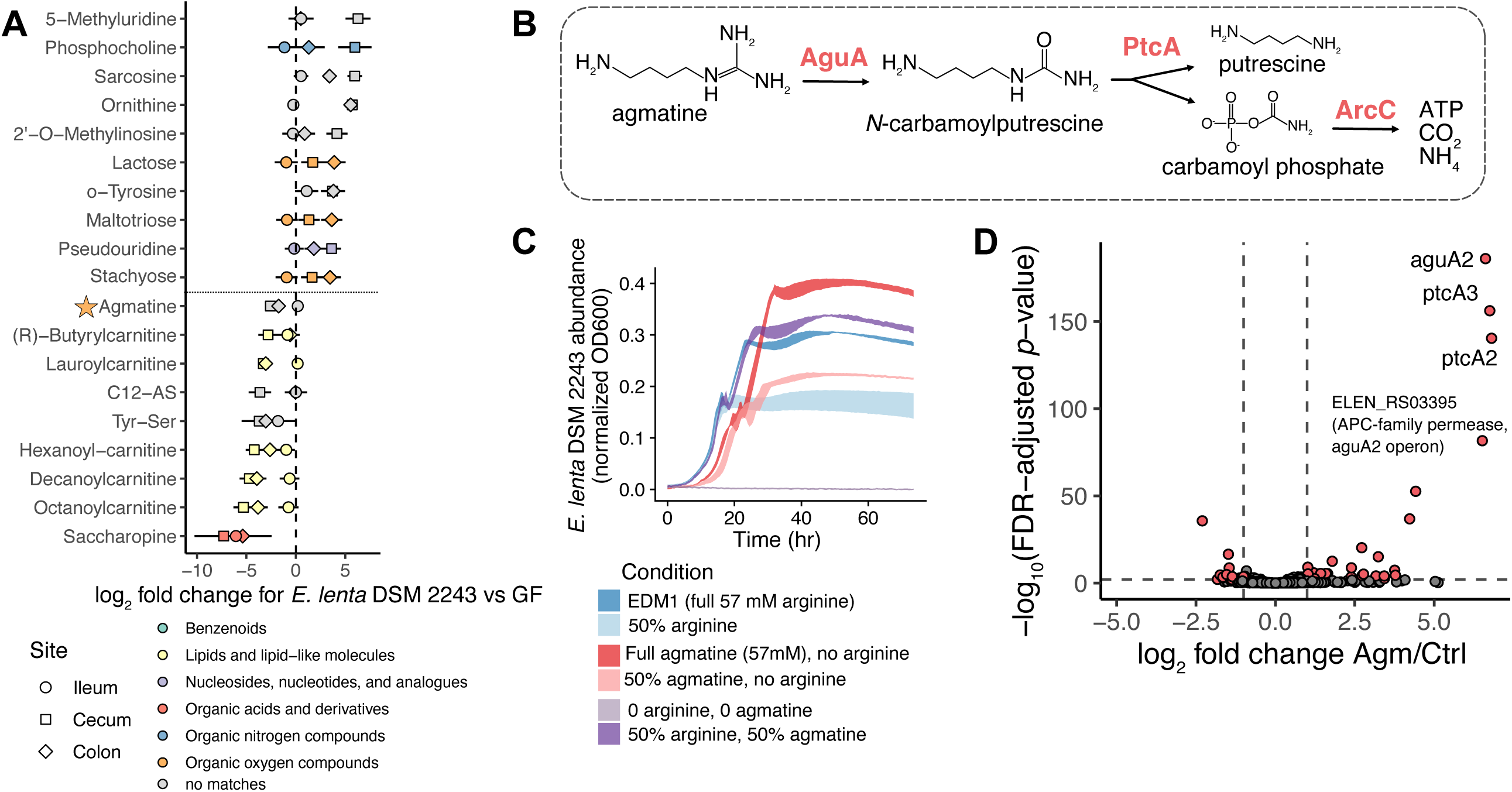
Agmatine can replace arginine as an energy source for *E. lenta.* **A)** Identified metabolite features with the highest estimated effects in *E. lenta* DSM 2243-colonized mice compared with germ-free. Each point indicates the effect size of that feature in a particular sample site (denoted by shape). B) Model of the agmatine deiminase ATP-generating pathway (Llácer et al., 2007). Three copies of an operon containing genes for all three of the labeled proteins are annotated in the *E. lenta* DSM 2243 genome. C) Growth of *E. lenta* DSM 2243 in EDM1 where arginine has been fully or partially replaced with agmatine sulfate. Curves show mean ± standard error for four replicates. D) Induction of the agmatine deiminase pathway in *E. lenta* DSM 2243 cultures in response to the addition of agmatine. The volcano plot shows the log_2_ fold change and FDR-adjusted *p*-values of agmatine-treated cultures compared to vehicle (as estimated by negative binomial differential abundance models with DESeq2). See also Figure S9, Table S7.

Given these results, we evaluated what other substrates may be used as carbon or energy sources by *E. lenta in vivo*. The metabolites most strongly depleted by the presence of *E. lenta* DSM 2243 in the intestinal tract included several fatty acids conjugated with carnitine as well as multiple other nitrogen-containing metabolites: saccharopine and agmatine (Figure 6A). We chose to investigate agmatine usage further for several reasons: its chemical similarity to arginine, the presence of known agmatine utilization genes in the *E. lenta* type strain genome, evidence of a consistent decrease across all three strain colonization groups (**Figure S9A**), and its multiple roles as a microbial metabolite and a host metabolite involved in regulation of cell division and neural signaling (Piletz et al., 2013). The *E. lenta* DSM 2243 genome contains two complete and two partial operons encoding genes for the agmatine deiminase pathway. This pathway operates analogously to the arginine deiminase pathway, with ATP production via carbamate kinase as the final step (Figure 6B). Despite this similarity, the agmatine deiminase enzyme family is highly structurally distinct from arginine deiminase (Llácer et al., 2007). Presence of this pathway is conserved across strains, as other *E. lenta* genomes contain anywhere between one and four copies of the key genes for agmatine deiminase and putrescine carbamoyltransferase (KEGG, **Table S7A**). Additionally, a transcriptional regulator found adjacent to this operon in some strains was previously associated with *E. lenta* competitive fitness *in vivo* (Bisanz et al., 2020).

Based on these observations, we predicted that *E. lenta* may be able to grow in the absence of arginine if it is supplied with agmatine as an alternative energy source. A flux balance analysis simulation of *E. lenta* in agmatine-based EDM1 predicted a somewhat reduced maximum growth rate (0.54 vs. 0.96 hr^-1^) in this condition, with arginine synthesized for protein via its annotated biosynthetic pathway from glutamate. Indeed, we found that replacing arginine with agmatine introduced a growth lag but resulted in a slightly higher final carrying capacity than the equivalent amount of arginine (Figure 6C). We additionally investigated agmatine-responsive genes using RNA-Seq. We grew *E. lenta* DSM 2243 in a formulation of EDM1 with 70% of the standard levels of arginine and acetate, treated cultures with either concentrated agmatine solution or water, and extracted RNA for sequencing. The genes most strongly induced by treatment with agmatine were two copies of putrescine carbamoyltransferase, one copy of agmatine deiminase, and a transporter in the same operon (Figure 6D and **Table S7B**). Genes in the second complete agmatine deiminase operon (ELEN_RS110[05-15]) were not differentially expressed, suggesting that the annotation of this second operon may be incorrect and/or may be involved in metabolism of a related compound. Interestingly, the most strongly downregulated genes were two transport-related genes adjacent to the energy-conserving hydrogenase (Ech) complex (ELEN_RS078[45-50]), one of which has structural homology to the arginine-ornithine antiporter found in the arginine deiminase operon (ELEN_RS09745, 27.6% identity). These results indicate that *E. lenta* can generate ATP from agmatine as a distinct alternative to arginine both *in vitro* and *in vivo* and has extensive genetic machinery to efficiently and specifically use each of these compounds.

## DISCUSSION

In this study, we used custom growth media and untargeted metabolomics to profile the metabolism of a poorly understood gut microbe at a systems level. Although *E. lenta* is found at > 50% prevalence in gut microbiota of North American adults (Koppel et al., 2018) and linked to acute and chronic disease (Alexander et al., 2021), very little is known about its core metabolic properties. We documented an unusual set of carbon sources, nutrient dependencies, and secreted metabolites, and incorporated these into a genome-scale metabolic model that accurately recapitulated pathway activity and response to new environments. We further identified core and strain-variable properties across a large collection of strain isolates. Finally, we evaluated the extent to which these *in vitro* and *in silico* findings can inform our understanding of the host-associated *in vivo* metabolic activity of this organism. This broad strategy uncovered several specific new findings on *E. lenta*’s role in the gut microbial ecosystem and its potential effects on human hosts.

We first analyzed *E. lenta*’s metabolic footprint in a sensitive chemically defined environment using untargeted metabolomics. The extent and variety of compounds produced by *E. lenta* across multiple growth phases is consistent with previous experimental and theoretical work on “costless” metabolite secretions by diverse microbes (Chodkowski and Shade, 2020; Dunphy et al., 2021; Pacheco et al., 2019). In particular, many nucleotides and nucleic acid intermediates are synthesized by *E. lenta* and secreted without any apparent cost to growth. Secretion of these broadly useful metabolites may contribute to a previously observed outsized impact of *E. lenta* on the composition of synthetic communities (Venturelli et al., 2018). Interestingly, several small molecules produced by *E. lenta* in EDM1 and in mice are known to impact host immune signaling, including indole-3-acetate (Roager and Licht, 2018) and inosine (Li et al., 2021; Mager et al., 2020). Notably, the relative level of production of these metabolites and others varied widely across *E. lenta* strain isolates. Teichoic acids, identified here as another strain-variable feature, are also key targets of host innate immunity, with differential responses depending on their composition (van Dalen et al., 2020). While much focus has been deservedly paid to individual specialized immunomodulatory transformations performed by *E. lenta* (Alexander et al., 2021; Paik et al., 2022), our results suggest that *E. lenta*’s effects on host immunity may be multifaceted.

We elucidated the roles of three common gut metabolites in the metabolic network of *E. lenta*: arginine, acetate, and agmatine. First, we confirmed that conversion of arginine to ornithine is a core property of the *E. lenta* species. Production of ornithine was the most consistent metabolic feature across strains and environments. Our stable isotope analysis indicated that ornithine is primarily an end product of growth and is relatively inaccessible as a carbon source for *E. lenta.* However, ornithine is a favorable carbon and/or energy source for numerous other gut microbes (Noronha et al., 2018), including as a substrate for Stickland metabolism by gut bacteria including *Clostridioides difficile* (Girinathan et al., 2021; Liu et al., 2022; Pruss et al., 2022). Therefore, production of ornithine by *E. lenta* may promote the growth of other proteolytic bacteria in the surrounding gut ecosystem.

We also found that the presence of acetate has a dramatic effect on *E. lenta* growth and metabolism *in vitro.* Acetate is a ubiquitous microbial metabolite in the mammalian gut that varies in concentration (van der Hee and Wells, 2021). Previous studies have speculated that *E. lenta* may produce acetate via autotrophic acetogenesis (Harris et al., 2018; Hylemon et al., 2018). While our study does not resolve the question of whether *E. lenta* has a functional acetogenic Wood-Ljungdahl pathway, we found that environmental acetate is an important biosynthetic precursor for *E. lenta*, incorporated partially via a distinctive bifurcated citric acid cycle (Amador-Noguez et al., 2010; Huynen et al., 1999). If *E. lenta* is in fact an acetate consumer *in vivo*, as we have observed *in vitro*, this role may have ecological consequences. For example, *E. lenta* may compete for cross-fed acetate with other gut microbes, including the abundant, health-linked members of the Firmicutes that metabolize acetate to butyrate at high rates (Duncan et al., 2002; Muñoz-Tamayo et al., 2011). However, while we did not identify any compound that can replace the role of acetate in *E. lenta*’s metabolic network, the observation that *E. lenta* can grow to high carrying capacities in rich media and in germ-free mice presumably lacking acetate indicates that other undetermined compounds may be able to serve as equivalent carbon sources.

Finally, we identified agmatine as an alternate energy source for *E. lenta in vivo.* Agmatine is a host metabolite with multiple roles as a neurotransmitter, regulator of nitric oxide synthesis, and regulator and precursor of polyamine metabolism (Piletz et al., 2013). Although agmatine can be synthesized at low levels by the host, particularly in the brain, the gastrointestinal tract is thought to be a major source of systemic agmatine (Haenisch et al., 2008)—sourced either directly from the diet and/or from microbial metabolism. Dietary sources of agmatine include a variety of plant and animal products, with the highest known levels in fermented foods and alcoholic beverages (Galgano et al., 2012). Altered agmatine levels have been associated with a range of diseases, including depression and diabetes (Piletz et al., 2013). Notably, reduced agmatine levels in the gut have been linked to cell proliferation and cancer (Molderings et al., 2004). Therefore, depletion of gastrointestinal agmatine by gut microbes including *E. lenta* has the potential to impact host health and disease. Further work is needed to clarify the roles of both production and degradation by gut microbes in regulation of host agmatine metabolism.

Overall, our analysis of *E. lenta* nutrient dependencies revealed that this species occupies a metabolic niche that is distinct from canonically described roles in the gut ecosystem, such as primary and secondary carbohydrate degraders or conventional methanogens and acetogens. *E. lenta* relies heavily on ATP generation from arginine and/or agmatine catabolism, uses acetate as a key carbon source, and likely performs anaerobic respiration with unknown and potentially diverse substrates. The carbon and energy sources and auxotrophies that we identified were highly conserved across the *E. lenta* species, with the exception of pantothenate. Knowledge of these conserved metabolic dependencies may be an important tool in future therapeutic attempts to engineer or modify *E. lenta* abundance, metabolic activity, and community interactions. In addition, the resources described here, together with the development of tools for genetic manipulation of *E. lenta*, may provide a basis for further investigation of the biochemical and physiological mechanisms underlying its distinctive metabolic strategy.

Another resource generated by this study is a curated constraint-based genome scale metabolic model of *E. lenta.* Constraint-based modeling is a promising approach for predicting community interactions and ecosystem engineering (Heinken et al., 2021a), but to date, community metabolic modeling tools have been difficult to validate and have generated relatively limited insights beyond what could be obtained with simpler annotation methods. Our analysis highlights the importance of phenotype-based curation of individual reconstructions. Specifically, the initial semi-curated AGORA model of *E. lenta* did not support any growth in EDM1 and lacked a complete version of the agmatine deiminase pathway. Yet analysis of the more fully curated reconstruction enabled us to confirm key reactions during growth with arginine and agmatine *in vitro*, identify gaps representing potential novel enzymes, and uncover strain differences in vitamin dependence. These results suggest that the quality and predictive power of community metabolic models of the gut microbiota could be greatly improved by systematic data generation and refinement of reconstructions for a metabolically diverse sample of common taxa. Comparisons with growth in defined media conditions, -omics data, and strain conservation can assist with model validation even when genetic tools are not available.

Our approach combining untargeted metabolomics, genome-driven media development, computational modeling, and gnotobiotic experiments may be a useful strategy for accelerating scientific understanding of the biology of other understudied microbes. Each of these model systems and data types produced a broadly useful resource that partially supported findings from the others while also revealing novel facets of *E. lenta* metabolism. Together, our study sheds light on the unusual metabolic profile of an important member of the human gut microbiota, establishes a foundation for future mechanistic studies of this organism, and demonstrates a generalizable multidisciplinary approach to decipher the metabolic strategies of understudied microbes.

## ACKNOWLEDGEMENTS

We thank the UCSF Gnotobiotics Core staff (Jessie Turnbaugh, Kimberly Ly) for animal care support and assistance. This work was supported by the National Institutes of Health (2R01HL122593; 1R01AT011117; 1R01DK114034 to P.J.T., F32GM140808 to C.N.). P.J.T. is a Chan Zuckerberg Biohub Investigator and held an Investigators in the Pathogenesis of Infectious Disease Award from the Burroughs Wellcome Fund. We acknowledge SeqCenter for assistance with RNA-Seq library preparation and sequencing.

## AUTHOR CONTRIBUTIONS

Conceptualization, C.N. and P.J.T.; Methodology, C.N., J.S., J.E.B., B.D., and P.J.T.; Software, C.N., A.H., and I.T.; Formal Analysis, C.N., J.S., K.T., A.H., Y.L., and B.D.; Investigation, C.N., J.S., J.E.B., V.E., M.A., Y.L., and B.D.; Data Curation, C.N., J.S., Y.L., D.D., and B.D; Writing - Original Draft, C.N.; Writing - Review & Editing, C.N., J.S., J.E.B., V.E., M.A., K.T., A.H., Y.L., D.D., I.T., B.D., and P.J.T.; Visualization, C.N.; Supervision, I.T., D.D., B.D., and P.J.T.

## STAR METHODS

### RESOURCE AVAILABILITY

#### Lead contact

Further information and requests for resources and reagents should be directed to the Lead Contact Peter Turnbaugh.

#### Materials availability

This study does not contain newly generated materials.

#### Data and code availability

RNA sequencing data has been deposited in NCBI GEO and are publicly available as of the date of publication. Metabolomics datasets have been deposited in Metabolomics Workbench and are publicly available as of the date of publication. Processed metabolomics datasets, growth data, and metabolic reconstructions are available from Zenodo and are publicly available as of the date of publication. Accession numbers and DOIs are listed in the key resources table.

All original code has been deposited at Zenodo and GitHub (https://github.com/turnbaughlab/2022_Noecker_ElentaMetabolism) and is publicly available as of the date of publication. DOIs are listed in the key resources table.

Any additional information required to reanalyze the data reported in this paper is available from the lead contact upon request.

### EXPERIMENTAL MODEL AND SUBJECT DETAILS

#### Mouse husbandry and experiments

Mouse samples analyzed in this study were collected and described previously in (Alexander et al., 2021). The mouse experiment was approved by the University of California San Francisco Institutional Animal Care and Use Committee. The mice were housed at temperatures ranging from 67-74°F and humidity ranging from 30-70% light/dark cycle 12hr/12hr. LabDiet 5021 chow was used. No mice were involved in previous procedures before experiments were performed. Mice were assigned to groups to achieve similar age distribution between groups.

C57BL/6J mice (males, ages 4-8 weeks) were obtained from the University of California, San Francisco Gnotobiotics core facility (gnotobiotics.ucsf.edu) and housed in Iso positive cages (Tecniplast). Mice were colonized via oral gavage with *E. lenta* monocultures (10^9^ CFU/mL, 200 μl gavage) and colonization was confirmed via anaerobic culturing and/or qPCR for an *E. lenta* specific marker (*elnmrk1*) (Bisanz et al., 2020; Koppel et al., 2018). Mice were colonized for 2 weeks prior to sacrifice and sample collection.

#### Bacterial strains

Strain isolates analyzed in this work are described in (Bisanz et al., 2020). All experiments were performed in an anaerobic chamber with 2-5% hydrogen gas, 20% carbon dioxide, and the balance nitrogen, with growth in a 37°C incubator. Standard BHI media (VWR 90003-040) supplemented with 1% L-arginine (referred to below as BHI+) was used for culturing outside of defined media experiments.

### METHOD DETAILS

#### Defined media formulations and preparation

Standard composition of the EDM1 media and related formulations are provided in **Table S1**. As specified in **Table S1**, some experiments were performed using the initial formulation of the media, and others using a simplified form based on the results of leave-one-out growth experiments. For most components, 30-1000x stock solutions were prepared following (Zhang et al., 2009). Stock solutions were sterilized with a 0.22 μm syringe filter and stored at −20°C. Amino acids were typically added together directly from powder into a combined 2x stock solution which was then filter sterilized with a 0.22 μm vacuum filter, except when preparing individual leave-one-out amino acid growth experiments. Most versions used ATCC Trace Mineral and Vitamin Mix Supplements (MD-TMS and MD-VS), except for experiments to test leaving out individual components of these mixes. Individual replacement components are specified in **Table S1**. Media formulations were allowed to equilibrate in an anaerobic chamber (Coy) for at least 24 hours prior to use.

#### Bacterial culture and growth assays

For growth and metabolomics experiments, glycerol stocks were first streaked on BHI+ agar plates and incubated at 37°C for 2-3 days. Individual colonies were inoculated into 3-4 mL liquid BHI+ and incubated at 37°C for 40-48 hours, or until approximately early stationary phase. Culture optical density (600 nm wavelength absorbance, OD600) was measured using a Hach DR1900 spectrophotometer. 1 mL samples of BHI starter cultures were then centrifuged at 1,568 rcf for 4 minutes in a microcentrifuge (ThermoScientific mySpin 12) in the anaerobic chamber and resuspended in 1 mL sterile phosphate-buffered saline (PBS). For leave-one-out experiments, the resuspended cells were washed by centrifuging and resuspending in PBS again. The resulting suspension was vortexed and diluted to an approximate OD600 of 0.1, and used as inoculum into defined experimental conditions.

Growth assays were performed in standard 96-well microplates (Corning) at 37°C with a microplate reader (Biotek Eon or PowerWave). 180 μL of defined media were pipetted into each well, followed by 20 μL of inoculum. All experiments included at least three sterile control wells for each condition, into which 20 μL of sterile PBS was pipetted to establish consistent background OD600 measurements. Replicate wells were distributed pseudorandomly across the plate to control for plate layout effects, and inoculated wells were always paired with an adjacent control well of the same condition. 3-6 replicates were included for each condition. Plates were sealed with a transparent Breathe-Easy sealing gas exchange membrane (RPI). Every 30 minutes, plates were shaken at medium speed for 40 seconds, after which OD600 readings were performed.

After large metabolomics and RNA-Seq experiments (see below), culture purity was checked by plating and 16S rRNA gene Sanger sequencing, using standard primers (8F AGAGTTTGATCCTGGCTCAG and 1542R AAGGAGGTGATCCAGCCGCA).

#### Sample collection for metabolomics

Time course experiments were conducted in tubes in the anaerobic chamber in a 37°C incubator. For all metabolomics experiments, three independent culture replicates were included for each condition, with an equal number of uninoculated control tubes. Starter cultures and inocula were prepared as described above for growth assays. 5mLs of defined media was added to VWR glass culture tubes (53283-800) with screw caps. The PBS-washed inoculum was added to culture tubes to obtain an approximate starting OD600 of 0.001. A preliminary growth assay was conducted to define time points spanning the exponential growth phase in the tested conditions. At each time point, OD600 measurements of all inoculated tubes were first measured using a Hach DR1900 spectrophotometer, with a paired control tube to normalize for the background. 100 μL from each tube were then transferred into a 96-well microplate, which was sealed and removed from the anaerobic chamber. Plates were centrifuged at 1,928 rcf at 4°C for 8 minutes, after which supernatants were collected into fresh polypropylene tubes or plates, sealed, and flash-frozen in liquid nitrogen.

Two time course experiments were carried out with stable isotope-labeled substrates. Experimental groups included conditions in which sodium acetate in the defined media was replaced with ^13^C_2_ labeled sodium acetate (Sigma-Aldrich 282014), along with a matched experimental group with the same concentration of unlabeled substrate. The same procedure was followed for the arginine labeling experiment, using ^13^C_6_ labeled L-arginine HCl (Sigma-Aldrich 643440).

For the comparative strain metabolomics experiment, 96-well polypropylene deep well plates were prepared with 800μL of fresh media in each well. Starter cultures and inocula for 29 isolates of *Eggerthella lenta* and 1 isolate of *Eggerthella sinensis* (Bisanz et al., 2020) were prepared as described above for growth assays, except without final dilution, and 80 μL was used to inoculate wells, leaving a blank well in between every culture well to prevent cross-contamination. After 72 hours, OD600 measurements were taken, plates were centrifuged, and supernatants were collected as described above.

#### Targeted quantification of acetate

A subset of unlabeled supernatant samples from the acetate labeling time course were shipped to Stanford University on dry ice for targeted quantification of acetate.

Samples (20 μL) were first mixed with an internal standard solution (30 μL; 1 mM phenylpropionate-d9) in a V-bottomed, poly(propylene), 96-well plate, and extracted by mixing with 3 sample volumes of extraction solution (75% acetonitrile:25% methanol). The plate was covered with a lid and centrifuged at 5,000 rcf for 15⃞min at 4⃞°C. Supernatant was collected for derivatization before subjecting to LC–MS analysis.

Samples were processed using a derivatization method targeting compounds containing a free carboxylic acid. Extracted samples were mixed with 3-nitrophenylhydrazine (NPH; 200⃞mM in 50% acetonitrile) and N-(3-dimethylaminopropyl)-N′-ethylcarbodiimide (120⃞mM in 6% pyridine) at a 2:1:1 ratio. The plate was sealed with a plastic sealing mat (Thermo Fisher Scientific, #AB-0566) and incubated at 40⃞°C, 600⃞rpm in a thermomixer for 60⃞min to derivatize the carboxylate-containing compounds. The reaction mixture was quenched with 0.02% formic acid in 10% acetonitrile:water before LC–MS.

Samples were injected via refrigerated autosampler into mobile phase and chromatographically separated by an Agilent 1290 Infinity II UPLC and detected using an Agilent 6545XT Q-TOF (quadrupole time of flight) mass spectrometer equipped with a dual jet stream electrospray ionization source, operating under extended dynamic range (1,700⃞m/z). Chromatographic separation was performed using an ACQUITY Bridged Ethylene Hybrid (BEH) C18 column 2.1 x 100 mm, 1.7-micron particle size, (Waters Corp. Milford, MA), using chromatographic conditions published elsewhere (Liu et al., 2022). MS1 spectra were collected in centroid mode, and peak assignments in samples were made based on comparisons of retention times and accurate masses from authentic standards using MassHunter Quantitative Analysis v.10.0 software from Agilent Technologies. Acetate was quantified from calibration curves constructed with acetate-d4 as a standard using isotope-dilution MS with phenylpropionate-d9 as the internal standard. Calibration curves were performed in a modified base form of EDM1 lacking amino acids and other carboxylic acids. A background level of 1.05mM of acetate was subtracted to obtain the final quantities.

A plate layout error for supernatant samples from time points 4-7 in this experiment was noted based on the resulting acetate concentrations and corrected across datasets.

#### Untargeted metabolomics

Bacterial culture supernatant and sterile media, used in culture, were thawed on wet ice. Once thawed, samples were homogenized by inversion five times. Extracellular culture supernatant samples were prepared as follows: 20 μL of culture supernatant were extracted using 80 μL of a chilled extraction solvent at −20°C (1:1 acetonitrile:methanol, 5% water containing stable isotope-labeled internal standards). Samples were homogenized via pipette action, incubated for 1 hour at −20°C, centrifuged at 4°C at 6000 rcf for 5 min. The supernatant was transferred to a new plate and immediately sealed and kept at 4°C prior to prompt analysis via LC-MS/MS.

Intestinal samples (colon, cecum, ileum) were prepared individually using a single protocol as follows. Samples were kept frozen on dry ice and massed to at least 10 mg. Four microliters of −20°C extraction solvent (2:2:1 methanol:acetonitrile:water + stable isotope labeled internal standards) were added per milligram of intestinal sample. Six to eight 1mm zirconia silica beads were added to each sample followed by prompt bead beating (15 Hz, for 10 minutes). Following a 1 hour incubation in the −20°C freezer, samples were centrifuged at 4°C at 18,407 rcf for 5 minutes. Supernatant was collected and stored at −20°C prior to centrifugal plate filtration (0.2 micron polyvinylidene difluoride (PVDF) Agilent Technologies, Santa Clara CA) at 4°C at 4,122 rcf for 3 min. Collection plate was sealed and maintained at 4°C prior to prompt analysis.

Serum samples were first thawed on wet ice. 20 μL of serum was extracted with 4 volumes of methanol, containing stable isotope labeled internal standards. Samples were homogenized by vortexing for 20 seconds and placed in a −20°C for 1 hour to maximize protein precipitation. After freezer incubation, samples were centrifuged at 4°C at 18,407 rcf for 5 minutes. Supernatant was removed and dried under vacuum via centrivap (Labconco Corp.). Dried samples were then resuspended in 30 μL of 80% acetonitrile in water containing exogenous standard CUDA at 60 ng/mL. Samples were maintained at 4°C prior to prompt analysis.

Within each analysis batch, a small amount of each sample was removed and combined to create multiple technical replicate ‘pools’ which were analyzed intermittently throughout the analysis. These pools were used as external standards to ensure instrument stability across the batch. Additionally, method blanks were created using LC-MS grade water in place of supernatant, sterile media, serum, or intestinal contents. These blanks were used to ensure that reported metabolites were not inadvertently added during sample preparation.

Samples, sterile media, pools, and blanks were promptly added to a Thermo Vanquish Autosampler at 4°C in a Vanquish UHPLC (Thermo Fisher Scientific, Waltham, MA). Chromatographic separation was performed using an ACQUITY Bridged Ethylene Hybrid (BEH) Amide column 2.1 x 150 mm, 1.7-micron particle size, (Waters Corp. Milford, MA), using chromatographic conditions published elsewhere (Lai et al., 2018). Samples were analyzed on a Thermo Q-Exactive HF orbitrap mass spectrometer operated utilizing data dependent acquisition of MS2. Data was acquired independently in positive and negative modes via subsequent injections.

#### SIRM metabolomics

Intracellular extract samples were prepared with the following procedure, which was optimized for lysis of thick gram-positive cell walls: 600 μL of culture was transferred to an Eppendorf tube in anaerobic conditions and subsequently centrifuged at 10,000rcf for three minutes at 4°C, after which the supernatant was removed and the samples were immediately flash frozen to quench metabolites. 300 μL of cold methanol was then added to each pellet, followed by sonication on ice for 5 minutes and then shaking at 4°C for 4-12 hours. Samples were then centrifuged at 4°C at 15,000 rcf for 8 minutes, after which 120 μL of supernatant was transferred to fresh tubes and stored at −80°C until analysis. Prior to analysis, intracellular samples were dried at room temperature via Centrivap Benchtop Concentrator (Labconco Corp.). Samples were re-suspended in 60 μL of a chilled solution of 1:1 methanol and acetonitrile, with 24% water at −20°C containing the internal standards CUDA and VAL-TYR-VAL each at 60 ng/mL. Samples were centrifuged at 4°C, 4,122 rcf for 5 minutes and the supernatant transferred to a vial and immediately capped for LC-MS analysis.

Extracellular supernatant extraction for SIRM metabolomics was performed as described above (Untargeted metabolomics section) with one modification. In SIRM samples, deuterated internal standards were replaced with CUDA and Val-Tyr-Val to enable untargeted enrichment analysis. LC-MS/MS analysis conditions for SIRM metabolomics were identical to those used for standard untargeted metabolomics.

#### Untargeted metabolomics data processing

Untargeted metabolomics datasets were processed using MS-DIAL version 4.60 (Tsugawa et al., 2015). Metabolite features with intensity not greater than 3-fold elevated in samples compared to mean blank intensity were removed. Annotations were assigned using both local (Han et al., 2021) and global (Mass Bank of North America) tandem mass spectral libraries. Annotation confidence scores were assigned based on Metabolomics Standards Initiative (MSI) best practices (Fiehn et al., 2007; Schymanski et al., 2014). Briefly; MSI level 1 denotes library matches of accurate mass (*m/z*), retention time (RT) and tandem mass spectra (MS2). MSI level 2 follows the same rules as MSI 1, but allows for partial matching of MS2 spectra - as is prone to occur when experimental spectra are convoluted. MSI level 3 denotes a high scoring and visually confirmed match of MS2 spectra. MSI level 4 is assigned when exact stereospecificity cannot be determined by MS2 and chromatographic separation. MSI level 4 is often assigned to sugars, lipids, and polyphenols. MSI levels 1 and 2 could only be assigned to metabolites in our local library, for which authentic standards have been analyzed in the same chromatographic conditions as the samples being annotated. Post processing was performed using MS-FLO (DeFelice et al., 2017) for removal of erroneous features.

Processed datasets were further analyzed using Feature-based Molecular Networking and MolNetEnhancer in the GNPS web platform (Djoumbou Feunang et al., 2016; Ernst et al., 2019; Nothias et al., 2020; Wang et al., 2016), which assigned ClassyFire chemical classes to features based on molecular networking, independently of whether they were assigned a library identity.

To merge positive and negative ionization mode datasets from the same samples, duplicate features across datasets were identified as those with an expected mass difference of less than 0.02, a retention time difference of less than 0.1, and a Pearson correlation across samples of at least 0.7. If one or both members of a pair of duplicate features were assigned an identification, the feature with lower (more confident) MSI score was retained in the merged dataset. Otherwise, the positive mode feature was retained. The other feature was removed for downstream analysis.

Prior to statistical analysis, initial untargeted metabolomics feature tables were filtered to remove features with a high coefficient of variation across replicate samples (> 50%) and to remove potential technical outlier samples where the total signal from all features differed from the assay median by > 50%. Log_10_-transformed intensities were used for most statistical analysis, with the exceptions of SIRM datasets and the comparative strains dataset (for which values were approximately normally distributed without transformation). A pseudocount equal to 0.25 times the minimum non-zero value was added to the peak intensities for each feature before log transformation. Heatmaps of metabolite abundances were generated using the *ComplexHeatmap* package (Gu et al., 2016).

#### SIRM data processing

Intra- and extracellular untargeted data generated from SIRM experiments was analyzed separately using *Compound Discoverer* version 3.3 (Thermo Scientific, Bremen, Germany). Samples treated with labeled compounds were always paired with matched samples treated with unlabeled compounds in order to correct for naturally occurring isotope abundances. Unlabeled samples were used for compound detection and formula assignment via isotope pattern-based prediction, spectral library matches, or mass lists matches. The isotope patterns and formulas from the sample files then served as a reference for the detection of potential isotopologues per compound in the labeled sample type.

Specifically, the workflow consisted of the following nodes in Compound Discoverer: *Input Files*→ *Select Spectra*→ *Align Retention Times (ChromAlign)*→ *Detect Compounds (Legacy)* → *Group Compounds*→ *Predict Compositions*→ *Search Mass Lists*→ *Search mzCloud*→ *Mark Background Compounds*→ *Assign Compound Annotations*→ *Analyze Labeled Compounds*→ *Descriptive Statistics*→ *Differential analysis*.

The default settings from the “Stable Isotope Labeling w Metabolika Pathways and ID using Online Databases” workflow were used, with the following modifications:

1. Detect Compounds (Legacy): General– Min.Peak Intensity: 10000; Ions: [M+H]+1 or [M-H]-1 for positive and negative mode experiments respectively.
2. Group Compounds: Peak Rating Filter– Peak Rating Threshold: 4; Number of Files: 3.
3. Search Mass Lists: Search Settings– Mass Lists: Combined Hilic Mass mzRT library; Use Retention Time: True; RT Tolerance: 0.3 min; Mass Tolerance: 5 ppm.
4. Search mzCloud: DDA Search– Match Factor Threshold: 85
5. Mark Background Compounds: General– Max. Sample/Blank: 3
6. Assign Compound Annotations: Data Sources– Data Source #1: mzCloud Search; Data Source #2: MassList Search; Data Source #3: Predicted Compositions.
7. Analyze Labeled Compounds: Pattern Analysis– Intensity Threshold [%]: 2

Positive and negative polarity files were analyzed initially as separate studies with the following study definitions: Study factors including strain, replicate, substrate concentration, sample type, and time point were assigned. Sample types were assigned as either sample (unlabeled), labeled, or blank. These study factors interfaced with several nodes to reduce undesirable features and maximize reporting of quality high intensity peaks with potential for accurate measurement of ^13^C incorporation.

Results were filtered for non-blank formula assignment and absence in background samples. The MSI levels and labeling status for persisting entries were manually inspected for each compound and annotated onboard via custom tags. MSI levels were assigned based on the criteria previously described to match MS-Dial output. The mass isotopologue distributions were plotted to ensure reproducibility between replicates of various time points and detect anomalous labeling trends. The absence of reported enrichment in control samples processed as labeled samples was verified. A minimum threshold of 3% combined enrichment across all isotopologues other than M+0 was applied. This threshold was necessary for less abundant peaks where the ^13^C natural isotopic abundance correction introduces uncertainty in the M+1 and M+2 isotopologues.

A specification of the full Compound Discoverer workflow is available at https://github.com/turnbaughlab/2022_Noecker_ElentaMetabolism.

#### RNA-Seq

*E. lenta* DSM 2243 glycerol stocks were plated on BHI+ and incubated anaerobically at 37°C for three days. A single colony was then inoculated into 5mLs of BHI+ liquid culture and incubated at 37°C for 48 hours. 1 mL of the resulting culture was then centrifuged, washed once, and resuspended in equal volume PBS; all in anaerobic conditions. 220 μL of this inoculum were transferred into culture flasks containing 20 mL of EDM1 (70% carbon source reduced version, see **Table S1**) to obtain a starting OD600 of 0.01. After 20 hours of growth (early or mid-exponential phase), these cultures were treated with an additional 1.9 mL of sterile water or filter-sterilized solution containing either L-arginine (to reach a final concentration of 86 mM), sodium acetate (final concentration of 14.5 mM), or agmatine sulfate (final concentration of 30 mM). After 18 more hours (late exponential phase), 7.5 mL of each culture was collected into 15 mL conical tubes containing 5 mL of Qiagen Bacterial RNA-Protect (#76506). Cultures were centrifuged at 2,800 rcf at 4°C for 10 minutes, after which the supernatant was carefully removed. Pellets were extracted directly using the Qiagen RNeasy Mini kit (#74104) with modifications for difficult-to-lyse Gram positive bacteria. Samples were maintained on ice throughout the protocol. Briefly, 200 μL of TE buffer containing lysozyme (15mg/mL, #L4919) and 20 μL of Qiagen Proteinase K (#19131) were added to each pellet, vortexed gently, and incubated at room temperature for 10 minutes with shaking on an Eppendorf ThermoMixer at 900 rpm. 700 μL of Buffer RLT was then added to each tube and vortexed, after which the full contents were transferred to MP Biomedical Lysing Matrix E tubes (#116914500) and disrupted mechanically in a BioSpec Mini-Beadbeater-96 for 50 seconds. After disruption, tubes were centrifuged for three minutes at 15,000rcf and 850 μL of supernatant was transferred to fresh tubes. 590 μL of 80% ethanol was added to each sample and mixed by pipetting, after which lysates were transferred to Qiagen RNeasy spin columns and washed, following the RNeasy Mini kit QuickStart protocol including a single on-column DNase digestion (DNase #79254). After purification, RNA was eluted twice into 30 μL of nuclease-free water. RNA integrity was checked using an Agilent TapeStation 4150 and stored at −80°C.

RNA library preparation and sequencing was performed by the Microbial Genome Sequencing Center/SeqCenter. Samples were DNase treated with Invitrogen DNase (RNAse free, #AM2222). Library preparation was performed using Illumina’s Stranded Total RNA Prep Ligation with Ribo-Zero Plus kit (#20040529) and 10bp IDT for Illumina indices. Supplementary oligonucleotide probes specific to *E. lenta* rRNA and other highly expressed noncoding RNAs were incorporated during Ribo-Zero depletion (**Table S10**). Sequencing was done on a NextSeq 2000 with 2×50bp reads. Demultiplexing, quality control, and adapter trimming was performed with bcl-convert v3.9.3.

Reads were trimmed and quality filtered using *fastp* v0.20.0 (Chen et al., 2018) with the following parameters: *--trim_poly_g --cut_front --cut_tail --cut_window_size 4 -- cut_mean_quality 20 --length_required 15*. The Hisat2 aligner v2.2.1 (Kim et al., 2019) was used to map reads to the *E. lenta* DSM 2243 reference genome, downloaded from NCBI RefSeq (GCF_000024265.1). Gene-level read counts were obtained using the corresponding NCBI annotations and the *featureCounts* function in the R package *Rsubread* v2.6.4 (Liao et al., 2019), with the minimum quality score set to 1.

#### Construction, curation, and analysis of metabolic reconstructions

Genome-scale metabolic reconstructions were created from genome sequences of 25 *E. lenta* strains (Bisanz et al., 2020) using the DEMETER pipeline (Heinken et al., 2020, 2021b). Briefly, DEMETER performs systematic refinement of a draft genome-scale reconstruction, in this case generated through KBase (Arkin et al., 2018). Based on manually gathered experimental data, gap-filling solutions that had been manually determined in a subset of reconstructions are propagated by DEMETER to newly reconstructed strains. Moreover, DEMETER ensures correct reconstruction structure through use of a curated reaction and metabolite database and removes futile cycles resulting in unrealistically high ATP production. A test suite ensures agreement with the input experimental data and verifies model features such as mass and charge balance and feasible ATP production. The *Eggerthella lenta* DSM 2243 reconstruction underwent additional refinement of reactions and gene annotations against manually performed comparative genomics analyses (Heinken et al., 2020).

Reconstructions were analyzed using the Cobra Toolbox version 3.0 (Heirendt et al., 2019) in Matlab version 2018b, with the IBM Cplex solver version 128. Defined media concentrations were mapped from compound names to BiGG metabolite IDs (King et al., 2016) and converted to cell uptake rates over the duration of *E. lenta*’s exponential growth phase in batch culture (**Table S1**) using the *concToCellRate* function in the Cobra Toolbox and an approximate cell dry weight of 3.3×10^-13^ g, calculated based on colony forming units and dry biomass quantification from two aliquots of a late-exponential phase EDM1 culture. Additional compounds detected in sterile culture media with high confidence based on untargeted metabolomics were included in the simulation media with a fixed maximum uptake rate of 1 mM/gDW/hr.

The collection of *E. lenta* strain reconstructions included two reconstructions of the type strain: the DSM 2243 reconstruction which had undergone additional comparative genomics curation with PubSeed (Overbeek et al., 2014), and a slightly smaller and less refined reconstruction included in the AGORA2 collection (Heinken et al., 2020) based on genome resequencing of the ATCC 25559 version of the type strain. Neither reconstruction initially displayed nonzero growth in EDM1 using flux balance analysis. In order to facilitate interpretation of FBA results and avoid excess gap-filled reactions, we used the simpler *E. lenta* ATCC 25559 type strain reconstruction as the basis for subsequent curation and analysis. We transferred reactions present in the DSM 2243 reconstruction into this version if they were supported by genome annotations from other sources (Prokka (Seemann, 2014), GapMind (Price et al., 2022)) and/or by experimental growth or metabolomics data. We also performed several additional custom curations. Transporters were added for metabolites identified with high confidence (Metabolomics Standards Initiative level 1) and detected as secreted or depleted with a log_2_ fold change greater than 2 in the stationary phase strain collection metabolomics dataset (Figure 4). Several pathways were also modified based on growth assay results and/or pathway annotation software (Price et al., 2022) and (Pascal Andreu et al., 2021)). Curations were checked for viable growth in EDM1 using flux balance analysis. Reconstructions for the other 23 strains were only curated to ensure growth in EDM1 and to allow import/export based on metabolomics data, but not based on genome analysis with GapMind (Price et al., 2022) or the results of leave-one-out growth experiments, since those were only performed using the type strain. A complete summary of all curation steps is found in **Table S3**.

Flux balance analysis (FBA), parsimonious flux balance analysis (pFBA), and flux variability analysis (FVA) were performed using the Cobra Toolbox functions *optimizeCbModel*, *minimizeModelFlux*, and *fastFVA,* respectively. Flux variability ranges are reported for 99% of the maximum growth rate.

Metabolite uptake and secretion ranges estimated by FVA were compared with the stationary phase strain collection metabolomics dataset (shown in Figure 4). To compare metabolite data with FVA estimates, identified metabolites were first mapped from InChIKey metabolite IDs to KEGG IDs using the CTS Convert utility (Wohlgemuth et al., 2010) implemented in the R package *webchem* (Szöcs et al., 2020). KEGG IDs were then mapped to BiGG IDs using the BiGG database (King et al., 2016) and manually checked for consistency with compound IDs in the AGORA models. For purposes of this analysis, metabolites were considered produced if they had a log_2_ fold change greater than 0.5 in supernatants from at least one of the three type strain isolates included in the experiment (*E. lenta* DSM 2243 - UCSF, *E. lenta* ATCC 25559, and *E. lenta* DSM 2243 - DSMZ), and depleted if the log_2_ fold change was less than − 0.5.

To compare gene expression values with model flux estimates, we first ran pFBA and FVA for the modified EDM1 condition used for RNA-Seq (with 70% of the standard levels of arginine and acetate). We obtained the set of genes linked to reactions in the iEL2243_2 reconstruction, using NCBI BLASTn to map genes between different sets of annotations. Genes linked to multiple reactions were counted multiple times for each reaction, and vice versa. Only genes linked to reactions in the original ATCC 25559 reconstruction were included.

Similarly, to compare reaction knock-out predictions with strain variation, genes linked to reactions in the original ATCC 25559 reconstruction were mapped to annotations used in a previous pan-genome analysis of 31 non-clonal *E. lenta* genomes (Bisanz et al., 2020). In this previous analysis, amino acid sequence families were clustered across genomes using ProteinOrtho (Lechner et al., 2011) with cutoffs of 60% identity and 80% coverage. This analysis was then used to determine the number of strains in which each gene family in the ATCC 25559 reconstruction was present, and compare that distribution with the effects predicted by knockout analysis of the unconstrained model.

#### Strain comparative metabolomics analysis

Metabolites were classified as strain-variably produced/depleted if they were differentially increased/decreased (FDR-adjusted *p*<0.1 and absolute log_2_ fold change>0.5) in supernatants from at least 1 isolate strain but fewer than 29 (of the 30 isolates included in this experiment).

The phylogenetic and comparative genomics analyses used in this study were previously reported, including a core gene phylogenetic tree (Phylophlan), gene family clustering across strains (ProteinOrtho) and Prokka and GhostKoala annotation of all genomes (Bisanz et al., 2020).

Procrustes analysis was performed using the R package *vegan* v2.6-2, with evaluation of significance using the *protest* function (Oksanen et al., 2022). The *E. sinensis* isolate was excluded from Procrustes analysis to avoid skewing the distribution of phylogenetic distances.

The gene-metabolite association analysis was performed as described previously (Bisanz et al., 2020), with different cutoffs for prioritization. Briefly, all observed patterns of gene family presence-absence (based on clusters of 60% identity and 80% coverage) were enumerated across the collection of genomes. Log-transformed metabolite intensities were then tested for association with each presence-absence pattern using Welch’s t-tests. Using an initial cutoff of an FDR-adjusted *p*-value of 10^-4^, 39.0% of metabolite features were significantly associated with a gene cluster by this method. To further restrict results to those features most likely to depend on the presence of a gene, we filtered gene-metabolite links using two additional separability criteria. First, the difference in median log_10_ metabolite values between strain samples with and without the gene was required to be at least 0.4. Secondly, the 10th percentile log_10_ metabolite value for strains with the gene was required to be at least 0.4 above the maximum value in controls, and the 90th percentile value for strains without the gene was required to be lower than that value. Finally, only the highest association for each metabolite feature was retained. This additional filtering resulted in the final table of 84 gene family-metabolite links. KEGG pathway enrichment analysis of the final gene set was performed using *clusterProfiler* v4.0.5 (Wu et al., 2021) with a *p*-value cutoff of 0.1.

#### Cross-dataset untargeted metabolomics analysis

As described above, untargeted metabolomics datasets from supernatant, mouse intestinal contents, and serum were collected using the same chromatography and mass spectrometry methods. Pairs of features were compared across these datasets and linked if they were within 0.007 *m/z*, 0.5 minutes retention time, and had a cosine similarity of at least 0.205 between their MS2 spectra for positive ionization mode and 0.251 for negative ionization mode. Features for which MS2 spectra were not collected were linked to other features within 0.001 *m/z* and 0.2 retention time. Linked feature pairs were also required to be annotated as the same adduct. These *m/z* and retention time thresholds were chosen based on examination of the distributions of pairwise differences between features. The cosine similarity cutoffs were chosen as the 99.5th percentile of cosine similarity between a large sample of unrelated feature pairs: specifically, all pairwise comparisons of two sets of 200 randomly sampled features with retention times differing by at least 1 and *m/z* differing by at least 0.01. This procedure was repeated separately for positive and negative ionization mode datasets. Under these criteria, only approximately 0.5% of linked features assigned an identity were linked to features with a conflicting identity. Linked pairs of features were merged into shared metabolite IDs that were carried forward for cross-dataset analysis and comparison.

### QUANTIFICATION AND STATISTICAL ANALYSIS

All statistical analyses were performed in R v4.1.1, with data visualizations generated using the *ggplot2* package (Wickham, 2016). Statistical tests, sample size and standard error are reported in the figures and figure legends. Benjamini-Hochberg false discovery rate (FDR) correction was used to adjust for multiple comparisons in all cases.

#### Untargeted metabolomics statistical analysis

Differential trajectories across time series datasets were assessed using spline models implemented in the *santaR* package (Wolfer, 2022). Differential abundance analysis between supernatant samples and sterile controls at the final time point was performed using Welch’s t-tests after checking assumptions of normality. In one case where cross-contamination of sterile control tubes occurred at later time points, features at those time points were compared with control samples from the last uncontaminated time point.

Differential abundance analysis for the comparative strains dataset was performed using linear models with each strain identity as a covariate. Differential abundance analysis for the gnotobiotic mouse intestinal dataset was performed using linear mixed models with the R package *lmerTest* (Kuznetsova et al., 2017), incorporating fixed effects for intestinal site, colonization group, and the interaction between them; and nested random effects for cage and animal. The *difflsmeans* function with Benjamini-Hochberg multiple hypothesis adjustment was used to evaluate the statistical significance of differences of each colonization group vs. germ-free under this model.

#### Statistical analysis of growth curves

Growth curves were normalized based on average time-matched readings from blank control wells. Normalized values were used to fit logistic growth models for each well using the R package *growthcurver* (Sprouffske and Wagner, 2016). Low-quality model fits (sigma > 0.1) were removed prior to calculation of summarized parameter values.

#### Targeted metabolomics statistical analysis

Differential abundance analysis was performed using a linear model with terms for time point, strain, and their interaction. Differences from controls under the resulting model were estimated using Dunnett’s method as implemented in the package *emmeans* v1.7.5 (Lenth, 2022).

#### RNA-Seq statistical analysis

Differential expression analysis was performed using negative binomial generalized linear models implemented in the *DESeq2* package v1.32.0 (Love et al., 2014).

### SUPPLEMENTAL INFORMATION TITLES AND LEGENDS

#### Supplemental Figures

**Figure S1. EDM1 chemically defined media supports robust growth of *Eggerthella lenta* and enables sensitive metabolomics profiling. Related to Figure 1. A)** Summary of the composition of EDM1 media. The number in parentheses indicates the number of specific compounds in each category. **B)** Growth of *E. lenta* DSM 2243 in two commonly used media conditions (Brain Heart Infusion supplemented with L-arginine, and ISP-2 media supplemented with L-arginine), compared with three initial defined media formulations. **C)** Comparison of total number of differentially abundant features and identified differentially abundant features in this experiment compared to previous metabolomics profiling of *E. lenta*. The combination of chemically defined culture media and untargeted metabolomics methods used in this experiment allowed for greater detection of metabolites produced by *E. lenta.* D) Metabolomics profiling of compounds known to be present in the chemically defined media formulation EDM1. 22 media compounds were detected, most of which were not significantly depleted in *E. lenta* cultures over time. E) Hierarchical clustering of metabolite trajectories reveals distinct growth phases. Scaled average metabolite intensities across time points during growth in EDM1 were hierarchically clustered with complete linkage and cut into discrete clusters with a height of 1.6, distinguishing early-, mid- and late-produced and depleted metabolites. Cluster order is arbitrary. Annotated metabolites are listed below each cluster along with their Metabolomics Standards Initiative confidence level. Colors indicate ClassyFire metabolite classes as assigned by GNPS. Only clusters with at least 1 identified metabolite and at least 5 total features are shown.

**Figure S2. Effects of individual media components on growth of *E. lenta* DSM 2243. Related to Figure 1. A)** Growth curves for *E. lenta* DSM2243 growth in EDM1 media with individual media components removed. Gray curves indicate growth in full EDM1 media in the same experiment. Curves are shown as mean +/- standard error. Blue text indicates the growth parameters with significantly different values with and without the compound (Wilcoxon rank-sum test, FDR-adjusted *p* < 0.2; *r* - growth rate *k* - carrying capacity, *tmid* - time to mid-exponential, *auc* - area under the empirical curve). **B)** Distribution of median effects of removal of all tested compounds on growth parameters estimated by a logistic model. The dotted line indicates the median parameter estimate for the full EDM1 media across all experiments. Parameters were fitted with a logistic model implemented by the R package *growthcurver*.

**Figure S3. Environmental acetate concentrations affect growth and metabolite production of three *E. lenta* strains. Related to Figure 2. A)** Targeted quantification of acetate depletion in E. lenta EDM1 cultures. Acetate was measured at 2-3 time points in supernatant samples from three *E. lenta* strains during growth in EDM1 as well as sterile controls. Quantification was performed using a method for derivatization of carboxylic acids with 3-nitrophenylhydrazine and N-(3-dimethylaminopropyl)-N′-ethylcarbodiimide followed by targeted LC-MS/MS. Error bars show mean +/- standard error. Linear models of acetate concentration versus strain and time point were inferred for each media group, and differences from controls under the resulting model were estimated using Dunnett’s method. * indicates *p*<0.05, *** indicates *p<*0.001. **B)** Growth of three *E. lenta* strain isolates in EDM1 with 0, 1, or 10 mM sodium acetate. Mean +/- standard error across three replicates is shown. **C)** Acetate-responsive metabolites in supernatants from *E. lenta* AB8n2 and *E. lenta* Valencia. Metabolites shown are those that were assigned an identification, were differentially abundant compared with sterile controls (FDR-adjusted *p*<0.2), and had significantly different trajectories over time in the presence vs absence of acetate in either strain (based on smoothing spline regression with the R package *santaR,* FDR-adjusted *p*<0.25). Values shown are scaled log-transformed peak heights. The number in parentheses indicates the Metabolomics Standards Initiative confidence level for each metabolite annotation (see *Methods*).

**Figure S4. Consistent incorporation of acetate across three *E. lenta* strains based on stable isotope-resolved metabolomics. Related to Figure 2. A)** Growth of *E. lenta* strains in EDM1 with varying levels of sodium acetate (either stable isotope-labeled ^13^C_2_ or unlabeled). Optical density measurements were taken and supernatant samples were collected at each indicated time point. Mean +/- standard error across three replicates is shown. **B)** Average trajectories of labeled extracellular metabolites in three different strains of *E. lenta*. Metabolites shown are those with > 50% and > 5×10^4^ average peak area from labeled isotopologues in at least one time point in the 10 mM labeled acetate group. For metabolites detected in both positive and negative ionization mode, only positive mode is shown. The value in parentheses indicates the Metabolomics Standards Initiative annotation confidence level for each metabolite. **C)** Labeled metabolites of known identity in intracellular extracts across three strains of *E. lenta* (data for DSM 2243 matches Figure 2E). Each panel shows the average mass isotopologue distribution across three replicates for a single metabolite in intracellular extracts from time point 5 (39 hours, late exponential phase). Metabolites are labeled with the compound name and Metabolomics Standards Initiative annotation confidence level in parentheses. Metabolites included are those with > 15% and > 10^4^ average peak area from labeled isotopologues in either the 1 mM or 10 mM labeled acetate group. *N*-acetylated amino acids are excluded for space and reported in **Data S1**. The isotopologue color legend is the same as in panel B. **D)** Labeled metabolites of unknown identity across three strains of *E. lenta*. Each panel shows the average mass isotopologue distribution (across three replicates) for a single metabolite in intracellular extracts from time point 5 (39 hours, late exponential phase). Metabolites are labeled with their estimated exact mass, retention time, and ionization mode. Metabolites included are those with > 15% and > 10^4^ average peak area from labeled isotopologues in either the 1 mM or 10 mM labeled acetate group. The isotopologue color legend is the same as in panels B and C.

**Figure S5. Stable isotope profiling of *E. lenta* arginine metabolism confirms that arginine is primarily converted to ornithine as an energy source. Related to Figure 2. A)** Citrulline, but not ornithine, has a similar effect as L-arginine on *E. lenta* growth. Growth curves of *E. lenta* grown in EDM1 media where the 1% L-arginine (red) has been replaced with an equimolar quantity of either L-citrulline (blue) or L-ornithine (green). Curves show mean +/- standard error across four replicates. **B)** In *E. lenta* DSM 2243 cultures grown with 1% ^13^C_6_ labeled arginine, correspondingly labeled citrulline and ornithine accumulate in supernatants over the course of growth. Curves show mean +/- standard error across three replicates. **C)** Mass isotopologue distributions of extracellular metabolites. Each barplot shows the isotopologue mean peak areas for each feature over time. Compounds shown are those of known identity that increase by a factor of at least 2^4^, have at least one isotopologue with a peak area of greater 10^6^ in at least one time point, and have a labeled isotopologue with >3% abundance in at least one time point. **D)** Mass isotopologue distributions of intracellular metabolites. Each barplot shows the mean peak areas of isotopologues for each feature at two time points. Compounds shown are those of known identity with an average labeled MID > 0.1 and a total peak area from labeled isotopologues of at least 10^5^ in at least one time point. The isotopologue color legend is the same as in panel C. **E)** Distribution of total signal of extracellular metabolites across labeling patterns. While signal from numerous unlabeled compounds is detected over time (left panel), compounds with M+5 labeling patterns are mainly restricted to ornithine, citrulline, and a compound of unknown identity (middle panel), and compounds found with high signal as M+6 isotopologues are mainly arginine and citrulline (right-hand panel). Compounds shown are those with the highest peak areas at the final time point in positive ionization mode. **F)** Hypothesized pathways for metabolism of L-arginine by *E. lenta*. Circles indicate the number of carbon atoms in selected compounds and are colored blue to indicate incorporation of ^13^C isotopes from external arginine. Compound names in bold were detected with the observed labeling patterns in either intracellular metabolite extracts or culture supernatants.

**Figure S6. Single-reaction knockout analysis of iEL2243_2 identifies conserved genes across metabolic subsystems. Related to Figure 3. A)** Predicted effects of knocking out reactions in the top 20 largest subsystems on growth of *E. lenta*, according to pFBA analysis of the iEL2243_2 model. Reactions designated “Has effect” are those for which the knockout has a predicted maximum growth rate less than wild-type but greater than 0. Essential reactions are those that reduced biomass flux to 0 when removed from the model. **B)** Reactions linked to more conserved gene families are more likely to have substantial effects on growth when removed. Each point represents a reaction, separated on the *x*-axis by whether the model without that reaction grew at > 70% of the wildtype model. The *y*-axis indicates the fraction of *E. lenta* strain genomes in which gene families (defined using ProteinOrtho clustering) linked to that reaction were present.

**Figure S7. Within-species variation in *E. lenta* metabolic profiles across genomes and metabolomes. Related to Figure 4. A)** Phylogeny of 30 *Eggerthella* strains analyzed in this study. This phylogeny was previously constructed based on core gene alignments using Phylophlan (Bisanz et al., 2020). **B)** Principal components analysis (PCA) of log-transformed metabolite intensity profiles of stationary phase supernatants from 30 *Eggerthella* isolates in EDM1. The right panel shows the largest feature loadings for the PCA and their corresponding chemical classes as assigned by GNPS, where available. Dereplicated metabolite features with an average value > 10^5^ in at least one strain were included. **C)** Distribution of the number of strains producing or depleting each metabolite feature. Features included are those that were significantly modified by at least one *Eggerthella* isolate in this experiment (FDR-adjusted *p*-value<0.1 and log_2_ fold change>0.5). **D)** Map of the teichoic acid biosynthesis region of the genome of representative *Eggerthella* strains. Genes outlined in bold are the gene families associated with the unidentified metabolite features shown in Figure 4E. Gene regions were defined in each genome based on the location of the genes annotated as *tagG* and *tagH* by Prokka. **E)** Distribution of core and accessory reactions across subsystems, based on comparative analysis of metabolic reconstructions of 24 *E. lenta* strain genomes. **F)** Predicted maximum growth rate inferred by flux balance analysis of each of the 24 *E. lenta* strain reconstructions in 52 leave-one-out media conditions based on EDM1. Gray tiles indicate predicted cases of zero growth.

**Figure S8. Differential abundance analysis of intestinal and serum metabolites of *E. lenta-*monocolonized mice compared to germ-free. Related to Figure 5. A)** Volcano plots of differential abundance analysis of metabolite features in intestinal contents and serum of gnotobiotic mice monocolonized with one of three *E. lenta* strains. Effect sizes and significance are estimated from group comparisons based on linear mixed models of log-transformed metabolite abundances, accounting for animal and cage random effects. **B)** Total number of untargeted metabolomics features in intestinal contents and serum of gnotobiotic mice that could be linked to features in either of two *in vitro* EDM1 metabolomics datasets, based on high similarity of *m/z*, retention time, and MS2 spectra. **C)** Comparison of the effect of *E. lenta* DSM 2243 on metabolites detected in both EDM1 cultures in the untargeted time course experiment and monocolonized mice. Each point represents a metabolite feature detected in both datasets. The *x-*axis indicates the log_2_ fold change of each feature in supernatants from the *E. lenta* DSM 2243 time course experiment compared with sterile controls, compared with the covariate-adjusted log_2_ fold change of that feature in monocolonized mice compared with germ-free mice. Points are colored green if the feature is significantly differentially abundant in gnotobiotic mice and is shifted in the same direction by the corresponding strain in the time course *in vitro* experiment.

**Figure S9. Shifts in intestinal amino acid metabolites of *E. lenta-*monocolonized mice compared to germ-free. Related to Figure 6. A)** Annotated metabolites with the largest shifts in intestinal contents of *E. lenta*-colonized mice compared with germ-free. Metabolites are shown if they were identified based on library comparison and were among the most 600 strongly shifted features in any individual site or colonization group, based on linear mixed models. Each point shows the effect size in a single site, and color indicates chemical class where available (assigned using feature-based molecular networking with GNPS). **B)** Abundance of arginine and agmatine-related metabolites in gnotobiotic mice. Arginine is only slightly depleted by *E. lenta*, although its expected products, ornithine and citrulline, are greatly increased. Agmatine is significantly depleted, while its expected product, putrescine, is not significantly increased. ‘.’ indicates Benjamini-Hochberg adjusted *p*<0.1, \**p<*0.05, \*\**p*<0.01, \*\*\**p<*0.001. **C)** Volcano plots illustrating shifts in the abundance of proteinogenic amino acids in *E. lenta*-colonized mice. Arginine is colored in green. Effect sizes and significance are estimated from group comparisons based on linear mixed models of log-transformed metabolite abundances, accounting for animal and cage random effects.

#### Supplemental Tables

**Table S1. Chemically defined media formulations used in this study. Related to Figure 1 and STAR Methods.** Recipes used for preparation of chemically defined media used for experiments in this study. The first two columns indicate the manufacturer information for each compound and the concentration of working solution prepared for that compound. Unless otherwise specified, reference to EDM1 indicates that the “Standard EDM1” preparation was used.

**Table S2. Summarized results of media leave-one-out growth experiments. Related to Figure 1.** Parameters were fit by logistic growth models using the R package *growthcurver.* A separate model was fit for each replicate in each experiment, and the average and standard deviation for each parameter across replicates are reported. Average growth rate *r* was calculated as a harmonic mean.

**Table S3. Curation steps applied to *E. lenta* DSM 2243 AGORA reconstruction. Related to Figure 3.** Summary of curation steps, supporting data, and gene annotations for each reaction added or modified in the iEL2243_2 reconstruction.

**Table S4. Most highly expressed genes by *E. lenta* DSM 2243 during growth in EDM1. Related to Figure 3.** Locus tags, gene annotation, and average and standard deviation of the 100 most highly expressed transcripts during *E. lenta* growth in the baseline EDM1 condition.

**Table S5. Metabolite features associated with variable *E. lenta* gene families across strains. Related to Figure 4.** Results of association analysis linking patterns of strain-variable genes with strain-variable metabolite features. Associations listed are those that met the strictest significance and separability criteria (see *Methods*). Gene annotations are listed for association patterns with 20 or fewer candidate gene families.

**Table S6. Summary of conserved and strain-variable reactions by subsystem in *E. lenta* strain metabolic reconstructions. Related to Figure 4.** Statistics on the distribution of core and accessory reactions across *E. lenta* strain metabolic reconstructions.

**Table S7. Genes linked to agmatine utilization by *E. lenta.* Related to Figure 6. A)** KEGG annotations of gene families in the agmatine deiminase pathway in *E. lenta* genomes, as previously obtained using GhostKoala. **B)** *E. lenta* DSM 2243 genes with differential expression in response to agmatine sulfate treatment (FDR-adjusted *p*<0.1 and absolute log2 fold change>1), as estimated by DESeq2.

**Table S8. Supplementary oligonucleotide probes used for depletion of highly abundant *E. lenta* noncoding RNAs. Related to STAR Methods.** Probes designed for depletion of *E. lenta* ribosomal RNA and highly abundant *ssrA* and *rnpB* noncoding RNAs, used in Illumina Ribo-Zero library preparation.

#### Supplemental Datasets

**Data S1. Labeled features detected in stable isotope experiments. Related to Figure 2**. Summary of labeled isotopologues detected by untargeted metabolomics. Each tab includes data for a single experiment and sample type: extracellular metabolites with labeled acetate, intracellular metabolites with labeled acetate, extracellular metabolites with labeled arginine, and intracellular metabolites with labeled arginine. In addition to basic properties of each compound/feature, the average peak area, standard error in peak area, and average fractional distribution are reported for each detected isotopologue. Compounds were filtered based on the same criteria as in Figures 2, **S6**, and **S7.**

**Data S2. Differentially abundant features across *in vivo* and *in vitro* untargeted metabolomics datasets. Related to Figure 5**. Each tab lists the set of untargeted metabolomics features that were differentially abundant (linear mixed effects models, absolute log_2_ fold change estimate > 1 and FDR-adjusted *p-*value < 0.2) in at least at least one intestinal site between *E. lenta*-colonized and GF mice, and that were also detected in *in vitro* untargeted metabolomics experiments, separated by strain and by feature annotation status (identified/unknown). For each feature, the corresponding log_2_ fold change and significance in the *in vitro* dataset(s) are listed for comparison. Features are ordered by their effect size in cecal contents.

