## Supplementary figures and images for "Systems biology illuminates alternative metabolic niches in the human gut microbiome"

### Figure S1

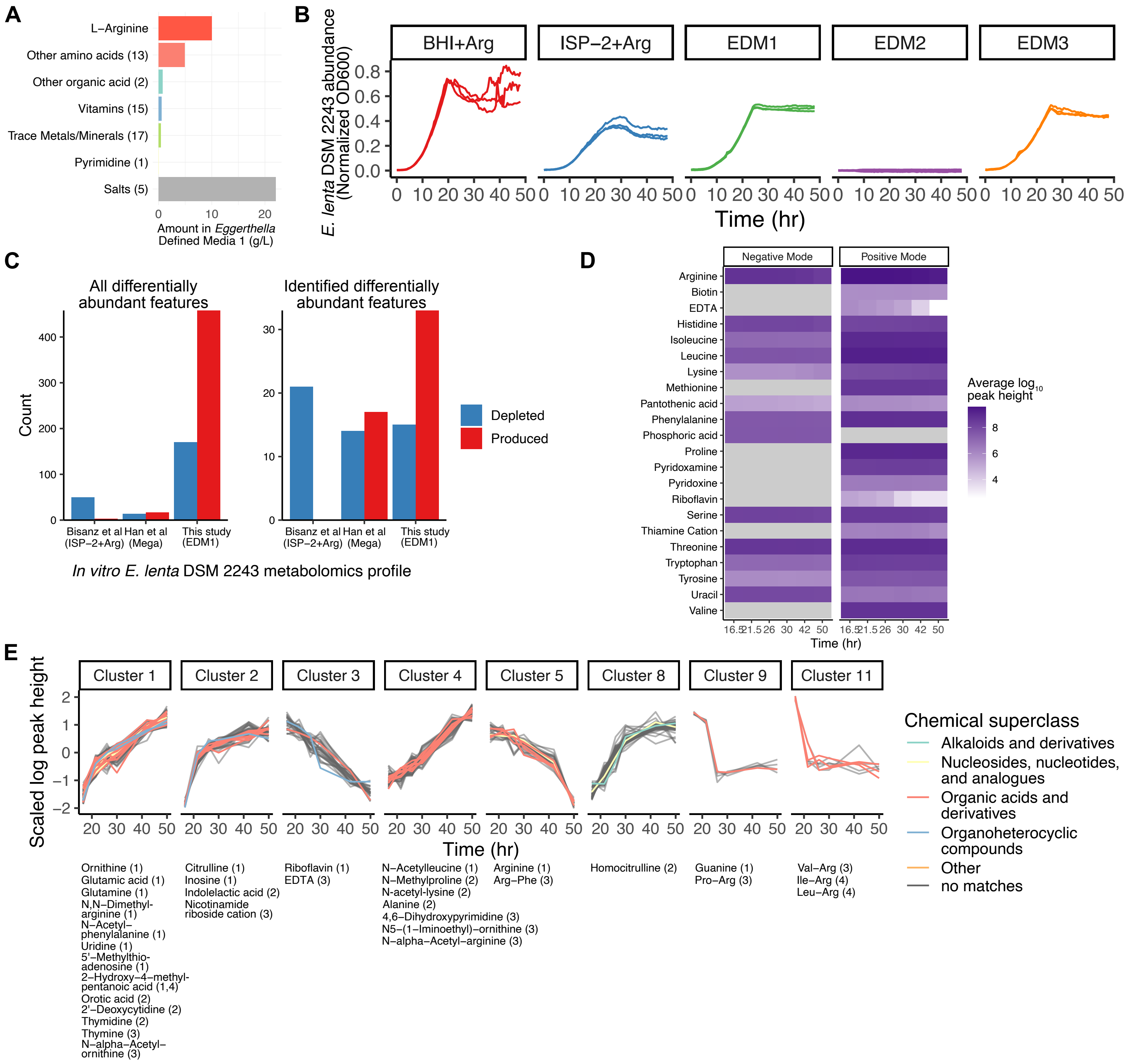

### Figure S2

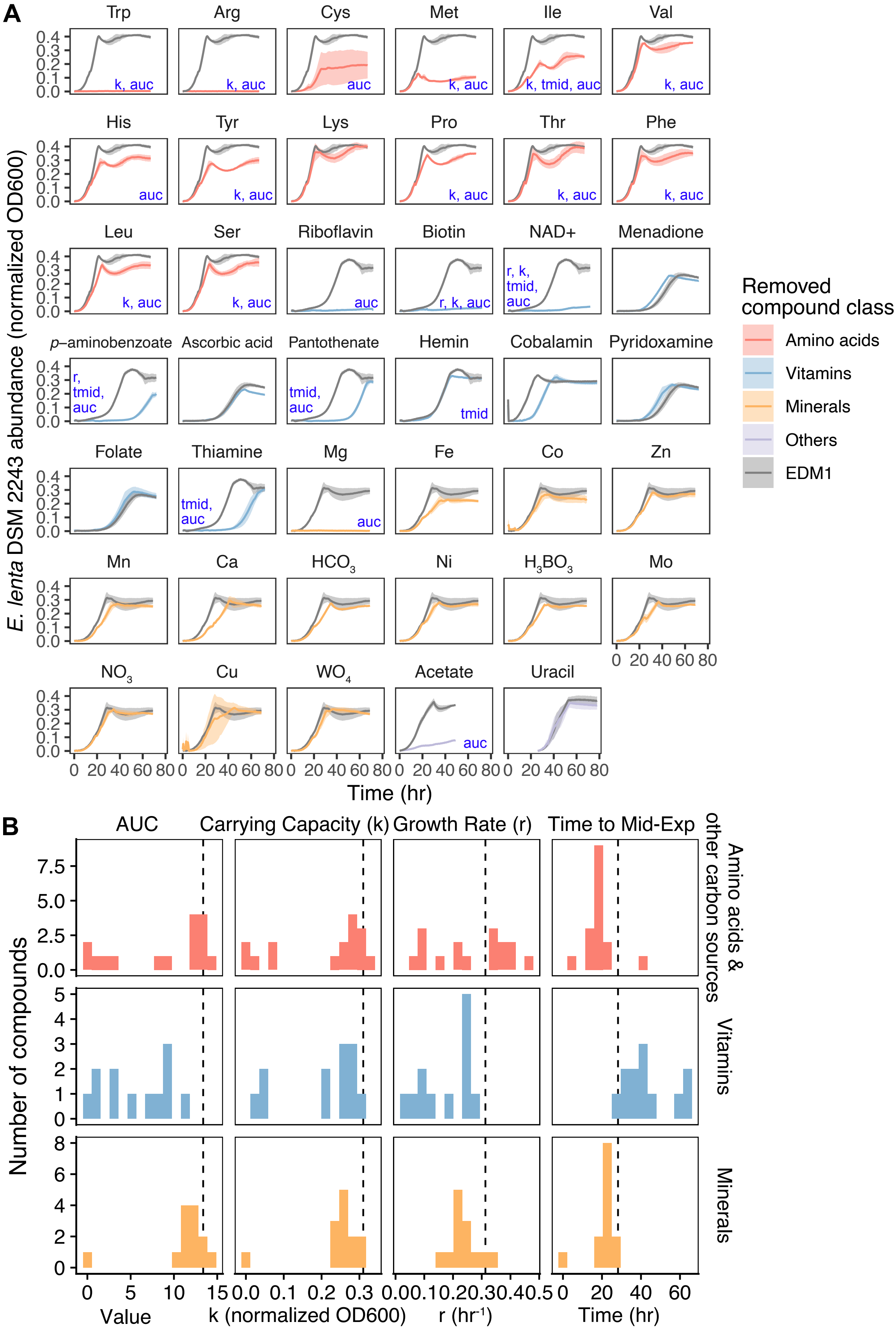

### Figure S3

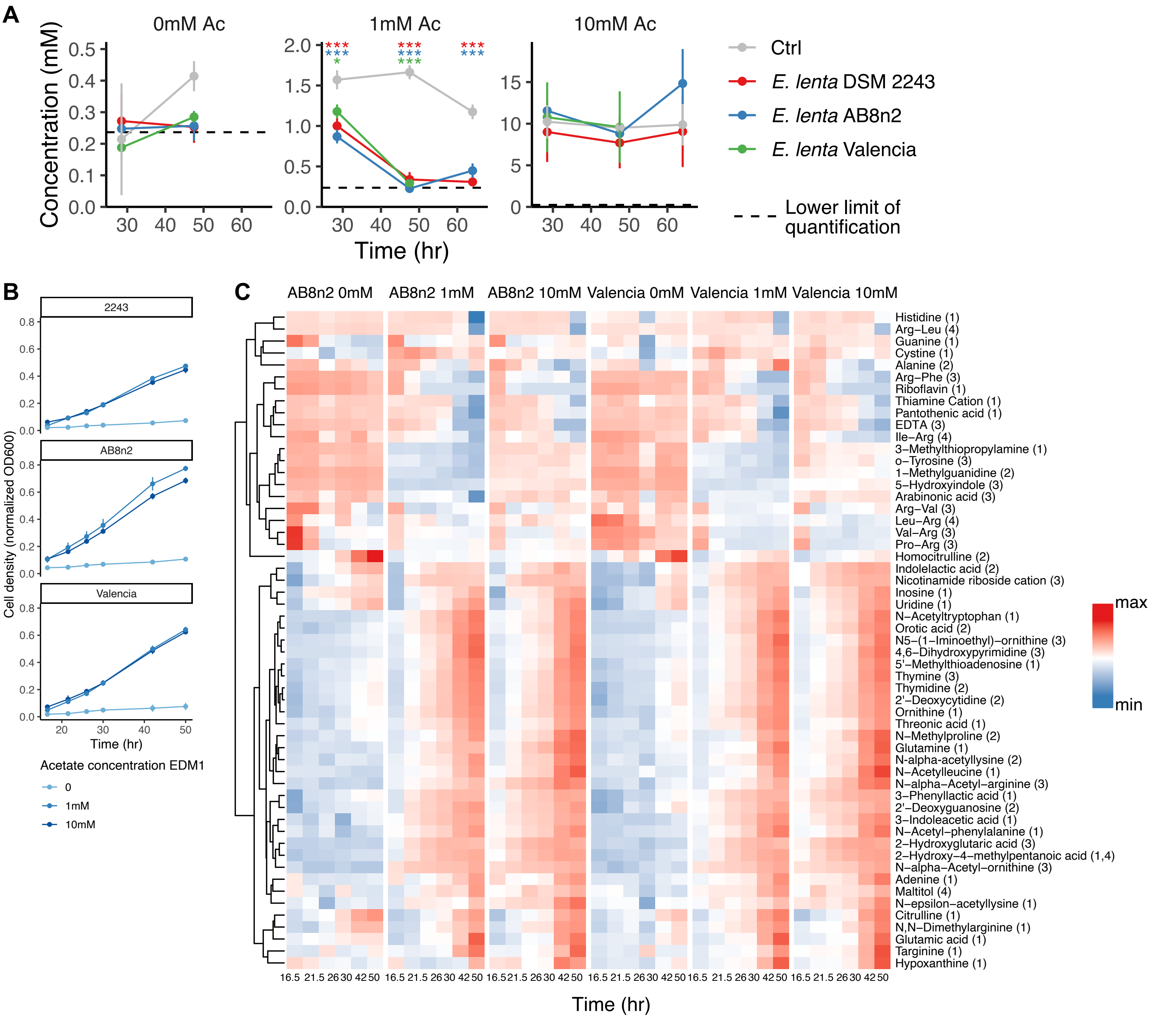

### Figure S4

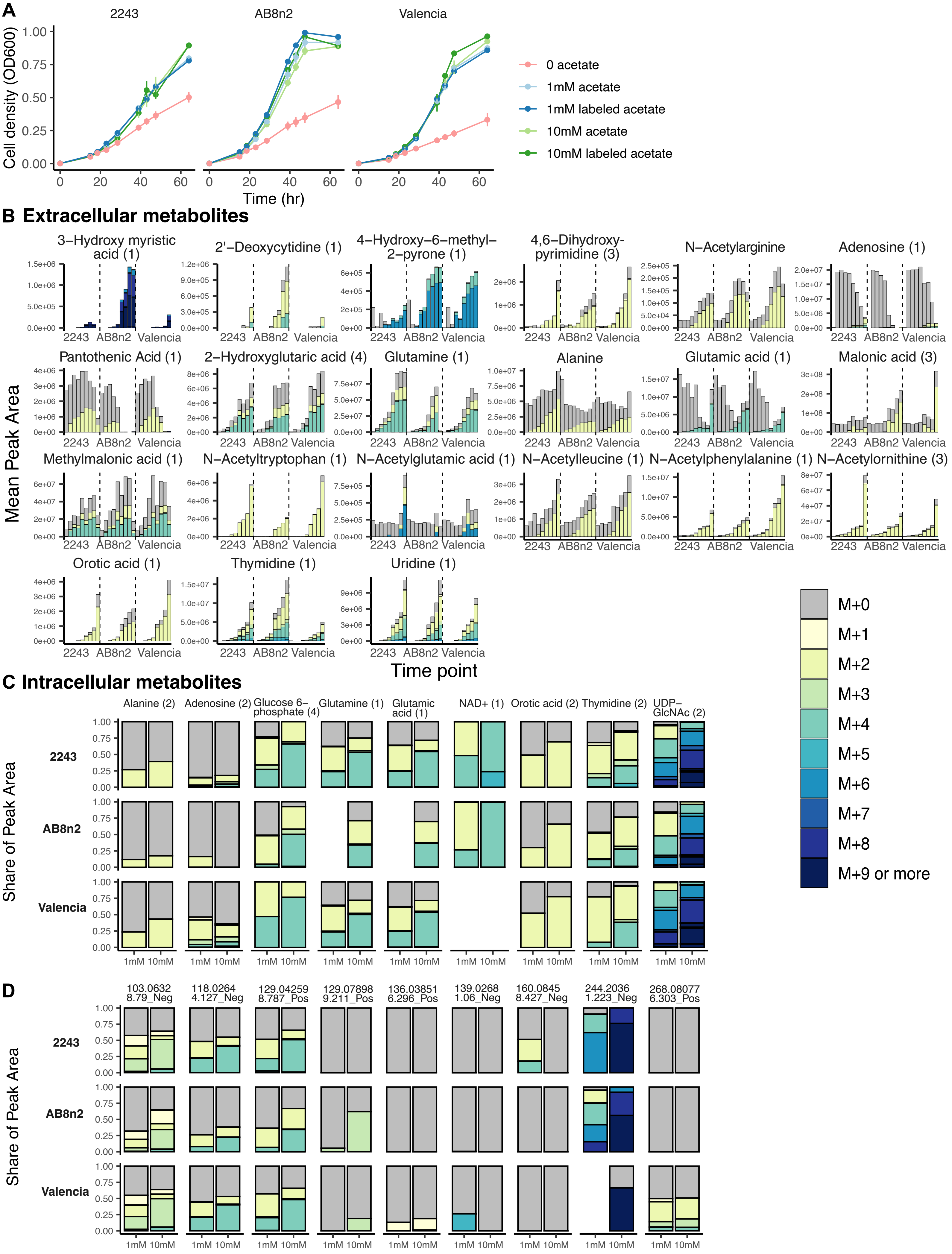

### Figure S5

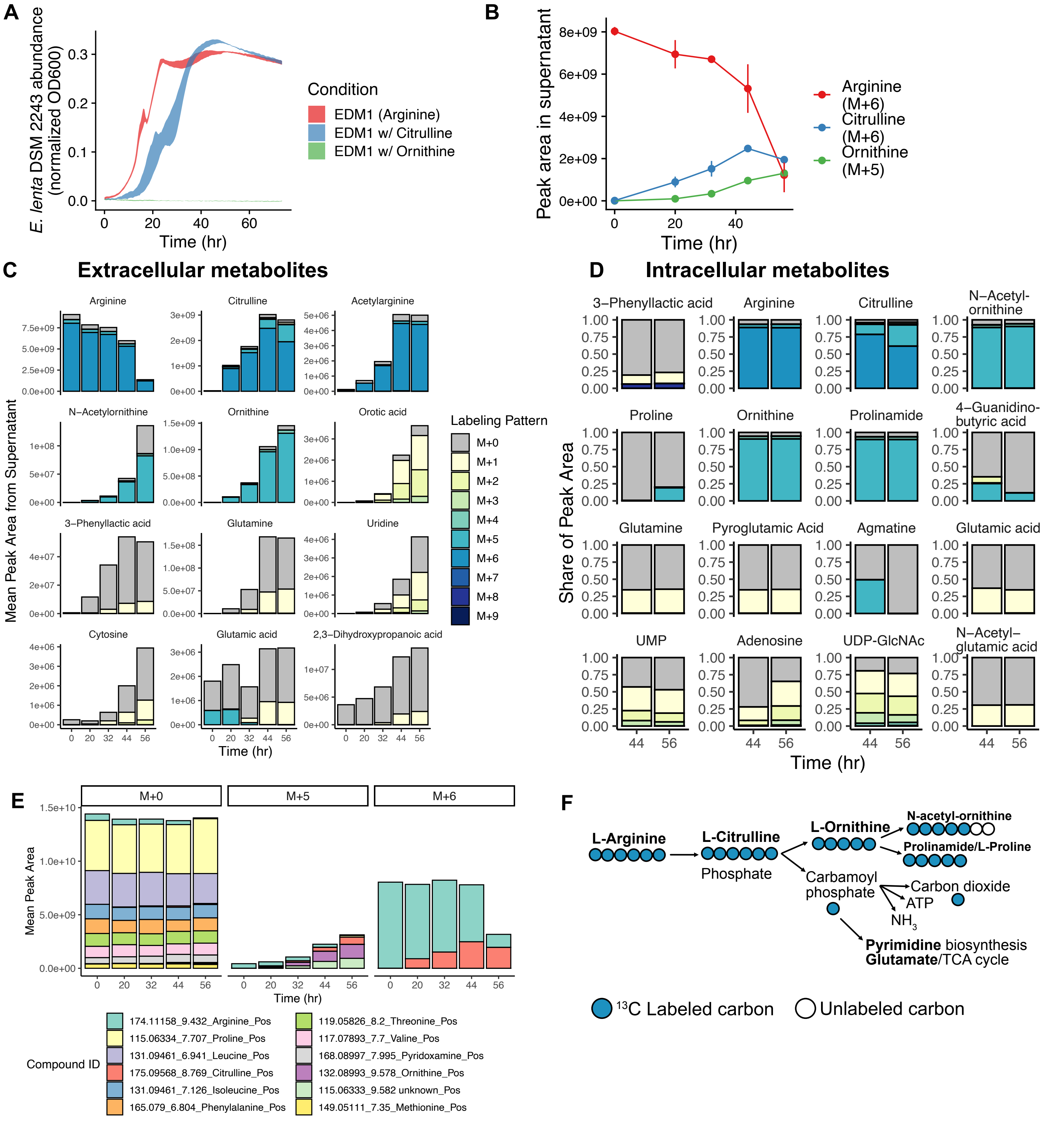

### Figure S6

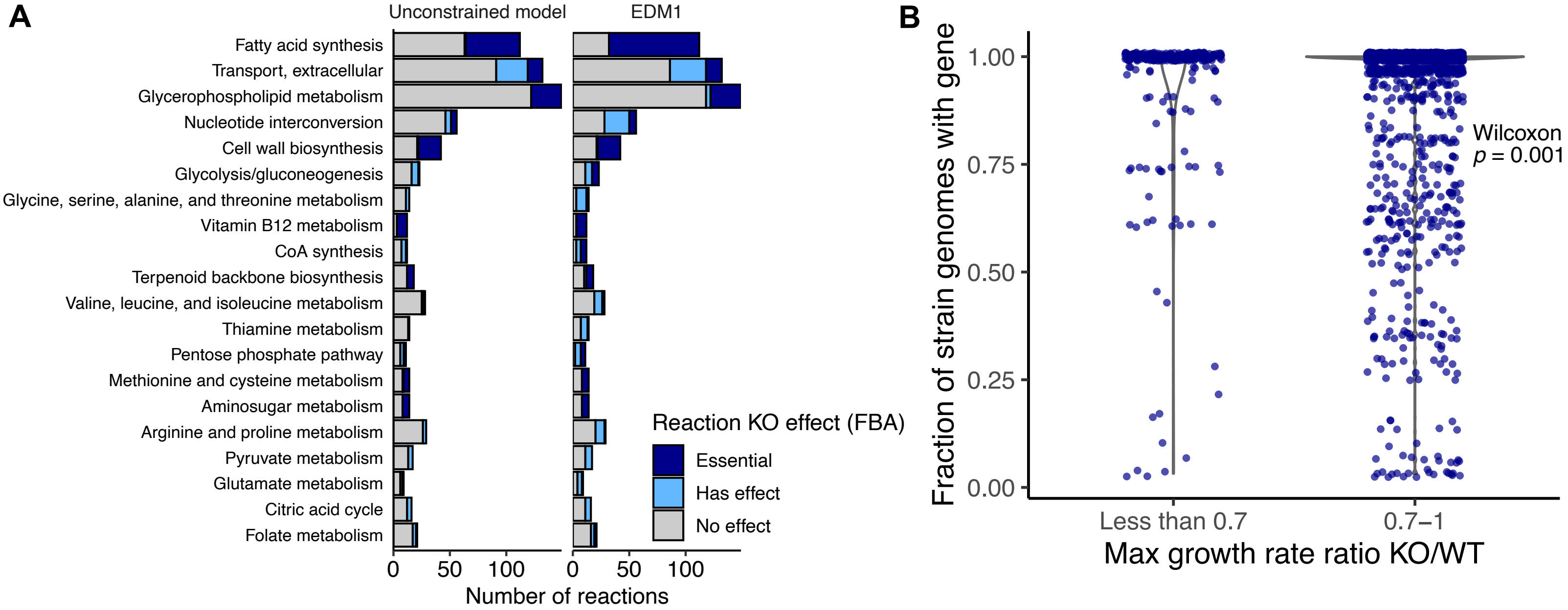

### Figure S7

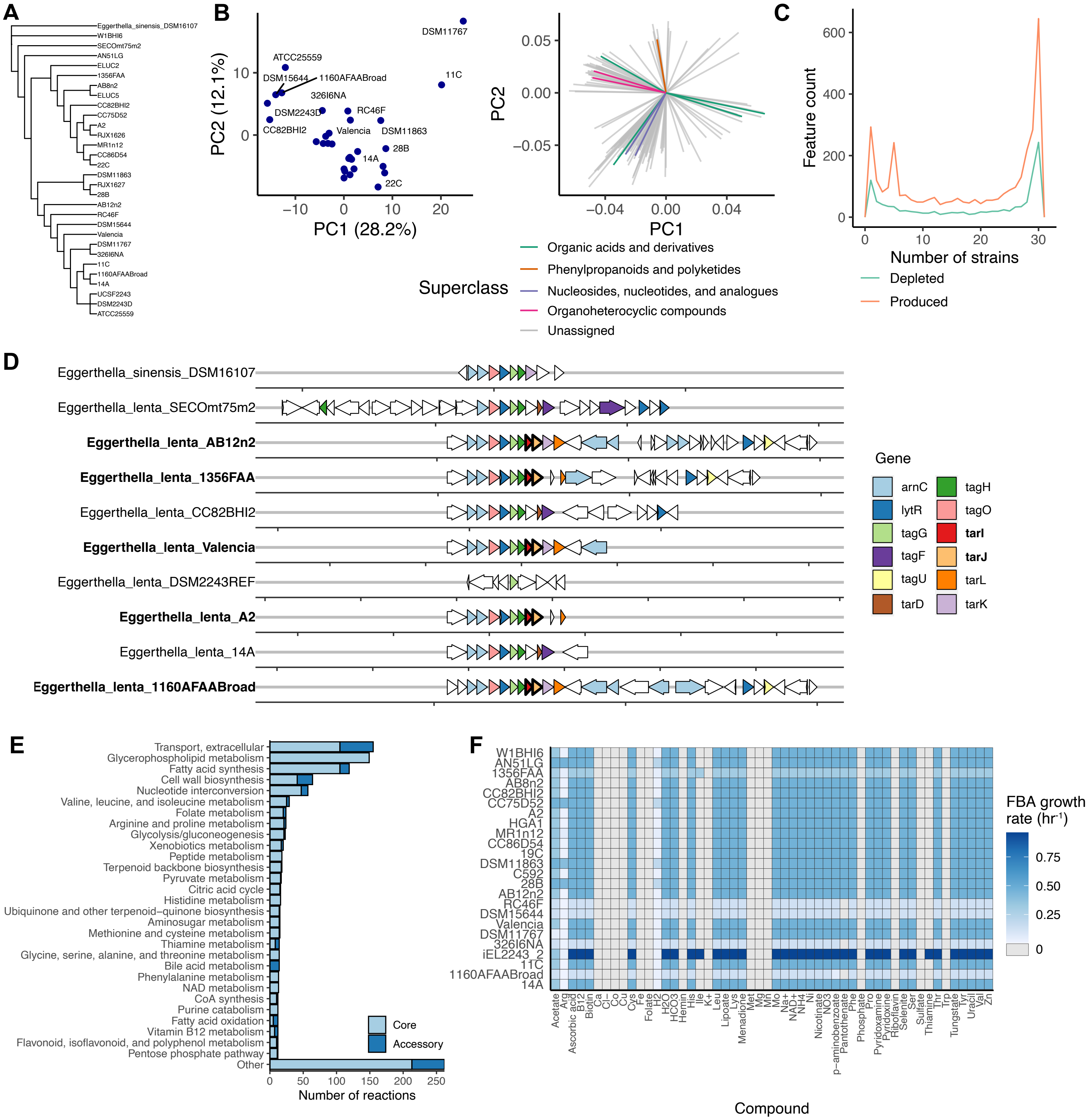

### Figure S8

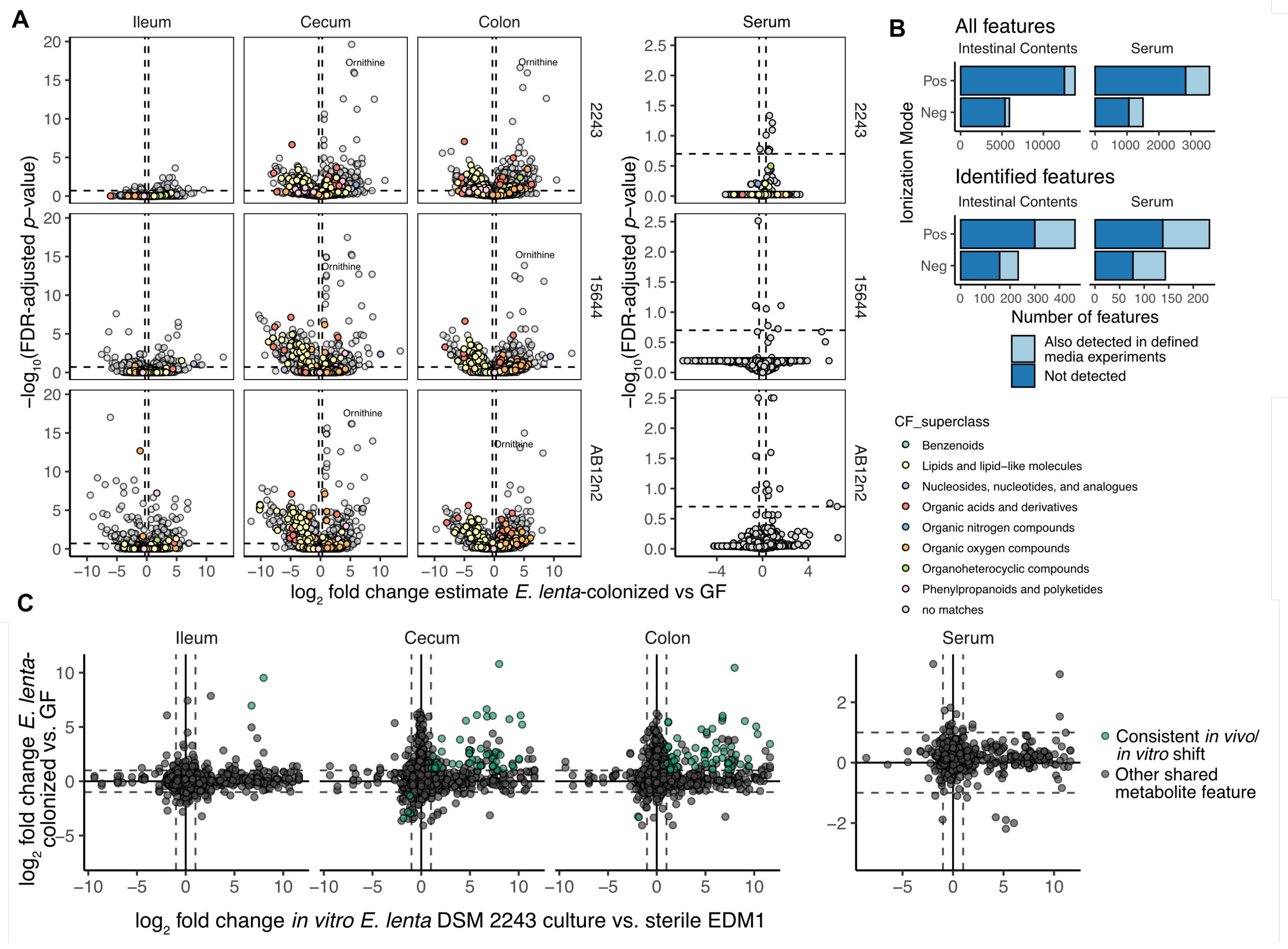

### Figure S9

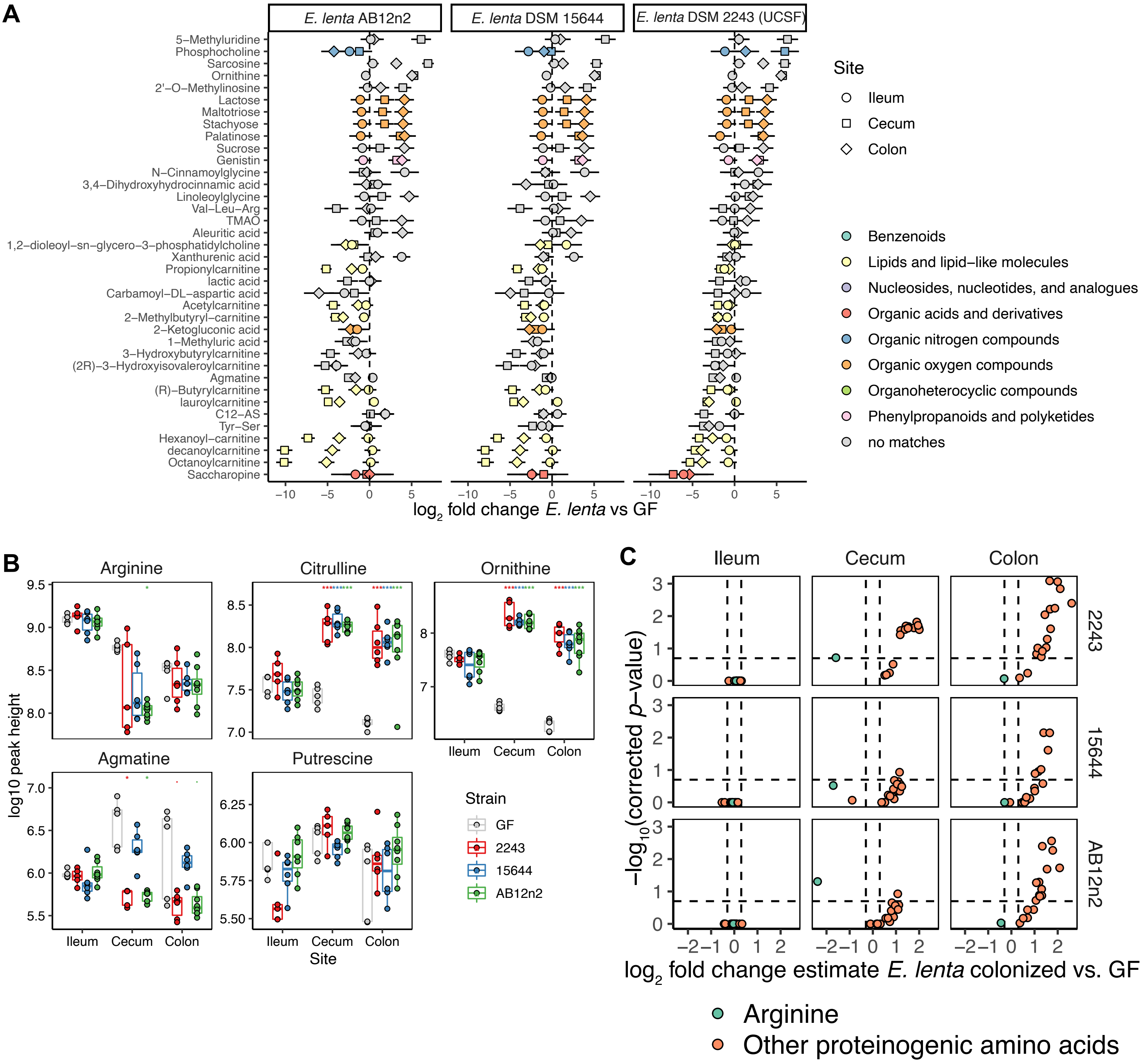
